# Evolutionary and functional data power search for obsessive-compulsive disorder genes

**DOI:** 10.1101/107193

**Authors:** Hyun Ji Noh, Ruqi Tang, Jason Flannick, Colm O’Dushlaine, Ross Swofford, Daniel Howrigan, Diane P. Genereux, Jeremy Johnson, Gerard van Grootheest, Edna Grünblatt, Erik Andersson, Diana R. Djurfeldt, Paresh D. Patel, Michele Koltookian, Christina Hultman, Michele T. Pato, Carlos N. Pato, Steven A. Rasmussen, Michael A. Jenike, Gregory L. Hanna, S. Evelyn Stewart, James A. Knowles, Stephan Ruhrmann, Hans-Jörgen Grabe, Michael Wagner, Christian Rück, Carol A. Mathews, Susanne Walitza, Daniëlle C. Cath, Guoping Feng, Elinor K. Karlsson, Kerstin Lindblad-Toh

**Affiliations:** Broad Institute of MIT and Harvard, Cambridge, MA, USA; McGovern Institute for Brain Research at MIT, Cambridge, MA, USA; Renji Hospital, School of Medicine, Shanghai Jiao Tong Univ., Shanghai, China; Program in Bioinformatics & Integrative Biology, UMass Medical School, Worcester, MA, USA; GGZ inGeest and Dept. Psychiatry, VU Univ. Medical Center, Amsterdam, the Netherlands; Department of Child & Adolescent Psychiatry and Psychotherapy, Psychiatric Hospital, Univ. of Zurich, Zurich, Switzerland; Neuroscience Center Zurich, Univ. of Zurich & ETH Zurich, Zurich, Switzerland; Zurich Center for Integrative Human Physiology, Univ. of Zurich, Zurich, Switzerland; Centre for Psychiatry Research, Department of Clinical Neuroscience, Karolinska Institutet SE-171 77 Stockholm, Sweden; Stockholm Health Care Services, Stockholm County Council, SE-14186 Stockholm, Sweden; Dept. Psychiatry, Univ. of Michigan, Ann Arbor, MI, USA; Dept. Medical Epidemiology & Biostatistics, Karolinska Institutet, Stockholm, Sweden; Dept. Psychiatry & Behavioral Sciences, USC, Los Angeles, CA, USA; Dept. Psychiatry & Human Behavior, Brown Medical School, Providence, RI, USA; Dept. Psychiatry, Harvard Medical School, Boston, MA, USA; BC Mental Health & Addictions Research Institute, UBC, Vancouver, BC, Canada; Dept. Psychiatry & Psychotherapy, Univ. of Cologne, Cologne, Germany; Dept. Psychiatry & Psychotherapy, Univ. of Medicine Greifswald, Greifswald, Germany; Dept. Psychiatry & Psychotherapy, Univ. of Bonn, Bonn, Germany; DZNE, German Center for Neurodegenerative Diseases, Bonn, Germany; Dept. Psychiatry and Genetics Institute, Univ. of Florida, Gainesville, FL, USA; Dept. Clinical & Health Psychology, Utrecht Univ., Utrecht, the Netherlands; Program in Molecular Medicine, UMass Medical School, Worcester, MA, USA; Science for Life Laboratory, IMBIM, Uppsala Univ., Uppsala, Sweden

## Abstract

Obsessive-compulsive disorder (OCD) is a severe psychiatric disorder linked to abnormalities in the cortico-striatal circuit and in glutamate signaling. We sequenced coding and regulatory elements for 608 genes implicated in OCD from humans and two animal models (mouse and dog). Using a new method, PolyStrat, which prioritizes variants disrupting evolutionarily conserved, functional regions, we found four strongly associated genes when comparing 592 cases to 560 controls. These results were validated in a second, larger cohort. *NRXN1* and *HTR2A* are enriched for coding variants altering postsynaptic protein-binding domains, while *CTTNBP2* (synapse maintenance) and *REEP3* (vesicle trafficking) are enriched for regulatory variants. The rare coding variant burden in *NRXN1* achieves genomewide significance (p=6.37×10^−11^) when we include public data for 33,370 controls. Of 17 regulatory variants identified in *CTTNBP2* and *REEP3*, we show that at least six alter transcription factor-DNA binding in human neuroblastoma cells. Our findings suggest synaptic adhesion as a key function in compulsive behaviors across three species, and demonstrate how combining targeted sequencing with functional annotations can identify potentially causative variants in both coding and noncoding regions, even when genomic data is limited.

## INTRODUCTION

OCD is a highly heritable (h^2^=0.27−0.65)^1^, debilitating neuropsychiatric disorder characterized by intrusive thoughts and time-consuming repetitive behaviors. More than 80 million people are estimated to be affected by OCD worldwide, and the majority do not find relief with available therapeutics^1^, underscoring the urgency to better understand the underlying biology. While specific genetic risk factors for OCD are unknown, genomewide association studies (GWAS) of OCD find an enrichment for genes in glutamate signaling and synaptic proteins^2,3^. In addition, past research in animal models, as well as genomewide genetic studies of disorders comorbid with OCD, identify genes and neural pathways likely to be involved in OCD.

In mouse, multiple genetically engineered lines have causally implicated the cortico-striatal neural pathway in compulsive behavior^4,5^. Mice with a deletion of *Sapap3*, which encodes a protein located postsynaptically in striatal neurons, exhibit self-mutilating compulsive grooming and dysfunctional cortico-striatal synaptic transmission, with abnormally high activity of medium spiny neurons (MSNs) in the striatum. The resulting compulsive grooming is ameliorated by selective serotonin reuptake inhibitor (SSRI), a first-line medication used to treat OCD^6^. Similarly, chronic optogenetic stimulation of the cortico-striatal pathway in normal mice leads to compulsive grooming accompanied by sustained increases in MSN activity^7^. Thus, excessive striatal activity, most likely due to a lack of inhibitory drive in MSN microcircuitry, is a key component of compulsive grooming. The brain region disrupted in this mouse model is also implicated in OCD by human imaging studies^8^.

Pet dogs are natural models for OCD amenable to genomewide mapping due to their unique population structure^9^, and genetic studies in dogs find genes linked to pathways involved in human compulsive disorders^10,11^. Canine compulsive disorder (canine CD) closely parallels OCD, with equivalent clinical metrics, including compulsive extensions of normal behaviors, typical onset at early social maturity, a roughly 50% rate of response to SSRIs, high heritability, and polygenic architecture^12^. Through GWAS and targeted sequencing in dog breeds with exceptionally high rates of canine CD, we associated genes involved in synaptic functioning and adhesion with CD, including neural cadherin (*CDH2*), catenin alpha2 (*CTNNA2*), ataxin-1 (*ATXN1*), and plasma glutamate carboxypeptidase (*PGCP*)^10,11^.

Additional genes are suggested by human genetic studies of related disorders such as autism spectrum disorders (ASD). Both ASD and OCD are characterized by compulsive-repetitive behaviors, and the disorders have high comorbidity (30-40% of ASD patients are diagnosed with OCD)^13–15^, suggesting a shared genetic basis^15^. Genomewide studies searching for both *de novo* and inherited risk variants have confidently associated hundreds of genes with ASD, and this set of genes may be enriched for genes involved in the compulsive-repetitive behaviors also seen in OCD^16^.

Using the research in mouse, dog and human, we can identify a large set of genes that is likely to be enriched for genes involved OCD. Focusing on this subset of genes in association studies could potentially accelerate the search for therapeutic targets by adding statistical power. Researchers, particularly in psychiatric genetics, are wary of studies that focus on sets of candidate genes, given that such associations often fail to replicate^17^. However, a closer examination of past studies suggests that this approach is both powerful and reliable when the set of genes tested is large, and when the study is looking for associations driven by rare variants^18^. For example, a large-scale candidate gene study testing 2000 genes for association with diabetic retinopathy identified 25 strongly associated genes, at least 11 of which also achieved genomewide significance in a GWAS of type 2 diabetes, a closely related disorder ^19,20^. Similarly, a recent targeted sequencing study testing 78 ASD candidate genes identified four genes with recurrent deleterious mutations; these four genes are also implicated by whole-exome sequencing studies^21^. Candidate gene studies also found replicable associations to rare variants in *APP*, *PSEN1* and *PSEN2* for Alzheimer’s disease^22^, *PCSK9* for low-density lipoprotein-cholesterol level^23,24^, and copy-number variants for autism and schizophrenia^16^.

A study design focused on a subset of the genome has the major advantage of allowing for a sequencing-based approach that, on a whole genome scale, would be cost-prohibitive. Sequencing-based association studies, unlike the more commonly used genotyping-based approach, capture nearly all variants in the regions targeted, facilitating identification of the actual causal variants underlying increased disease risk and accelerating the discovery of new therapeutic avenues^25–27^. For example, finding functional, rare variants in *PCSK9* led to new therapies for hypercholesterolemia^23,24,28,29^. Unbiased whole genome sequencing, which provides the most complete genetic information, is still prohibitively expensive at the cohort sizes required for complex diseases. Lower-cost whole-exome approaches, which target predominantly coding regions, have identified causal variants for rare diseases^30^, but do not capture the majority of disease-associated variants, which are predicted to be regulatory^31,32^. A targeted sequencing approach that captures both the regulatory and coding variation of a large set of candidate genes offers many of the advantages of whole genome sequencing, but is feasible when cohort size and resources are limited.

Here we report a new strategy that overcomes the limitations of less comprehensive candidate gene studies and exome-only approaches, and identifies functional variants associated with an increased risk of OCD. Drawing on findings from studies of compulsive behavior in dogs and mice, as well as studies of ASD and OCD in humans, we compiled a large set of 608 genes. We targeted both coding and regulatory regions in these genes for pooled sequencing in 592 OCD cases and 560 population matched controls, and analyzed the sequence data using a new method, PolyStrat, that incorporates regulatory and evolutionary information to find genes with an excess of functional variants in cases. We validated candidate variants by genotyping in an independent cohort of individuals. For coding variants, we additionally compared our data to the 33,370 individuals from the Exome Aggregation Consortium (ExAC)^33^. Finally, we directly tested our top candidate variants for regulatory function using electrophoretic mobility shift assays (EMSA).

Our results provide new insight into the biology of OCD, and may help to identify new therapeutic targets for this hard-to-treat disease. More broadly, our approach is applicable to genomic studies of any complex, heritable disease for which candidate genes and pathways can be identified from earlier studies, and is especially useful for investigating disease genetics in populations where assembling and sequencing large sample cohorts is not feasible.

## RESULTS

### Targeted sequencing design

We compiled, from multiple sources, a list of 608 genes previously implicated in OCD or related diseases. These genes fall into three categories, with 65 belong to more than one category(Table S1):

1. 263 ‘model organism genes’. These include 56 genes from regions associated in our GWAS of canine CD^10,11^ as well as 222 genes implicated in murine compulsive grooming, *i.e.* 68 genes encoding proteins differentially expressed in the MSN microcircuitry^34^ and 154 genes encoding proteins postsynaptically expressed^35–37^ in striatal neurons within the cortico-striatal pathway.
2. 196 ‘ASD genes’ implicated through animal models and whole genome studies of ASD in humans. ASD is comorbid with OCD and shares repetitive behavioral characteristics^38^.
3. 216 ‘human candidate genes’. These include 56 genes implicated by small-scale human candidate gene studies of OCD, 91 falling in chromosomal regions implicated by family-based linkage studies of OCD, and 69 from loci implicated in other neuropsychiatric disorders^39–45^.

We targeted the coding regions and 82,723 evolutionarily constrained elements in and around these genes for sequencing, comprising a total of 13.2Mb (58 bp-16 kb size range, median size 237 bp), with 34% noncoding^46^.

### Variant detection

We sequenced 592 European-ancestry DSM-IV OCD cases and 560 ancestry-matched controls using a pooled-sequencing strategy, with 16 samples per barcoded pool (37 “case” pools; 35 “control” pools). 95% of target regions were sequenced at >30x read depth per pool, with a median of 112x (~7x per individual; Figure S1), sufficient to identify variants occurring in just one individual, assuming a 0.5-1% per base machine error rate.

We detected variants in each pool using two algorithms: Syzygy^25^ and SNVer^47^. A total of 41,504 ‘high-confidence’ single nucleotide polymorphisms (SNPs) were detected by both, with highly correlated allele frequencies (AF) (ρ=0.999, p<2.2 × 10^−16^; Figure S2). In addition, 83,037 ‘low-confidence’ SNPs were detected by only one method (Syzygy: 42,712; SNVer: 40,325). The number and AF of high-confidence variants did not differ significantly between case and control pools, indicating no bias in variant detection. We used high-confidence variants for the primary analyses.

### Variant annotation

We combined evolutionary sequence conservation and biochemical annotations of DNase1 hypersensitivity sites (DHS) to identify likely functional variants in noncoding targeted regions. Regions conserved across species are exceptionally enriched for disease-associated variants^32^. DHS are central to cis-regulatory elements and delineate regulatory regions^48^, and variants in DHS sites explain a large proportion of complex disease heritability^31^.

We annotated variants as coding, DHS, and/or evolutionary (conserved or divergent)^49–51^. Overall, we annotated 67% (27,626) of high-confidence variants, of which 16% were coding (49% of those were nonsynonymous), 36% were in DHS, and 80% were evolutionary constraint or divergent (Figure 1a). For evolutionary annotations, we used the mammalian constraint GERP++ score^52^, where zero indicates no selective pressure. We annotate bases with GERP++>2 as ‘conserved’, bases with GERP++<–2 as ‘divergent’; and both as ‘evolutionary’.

**Figure 1.**
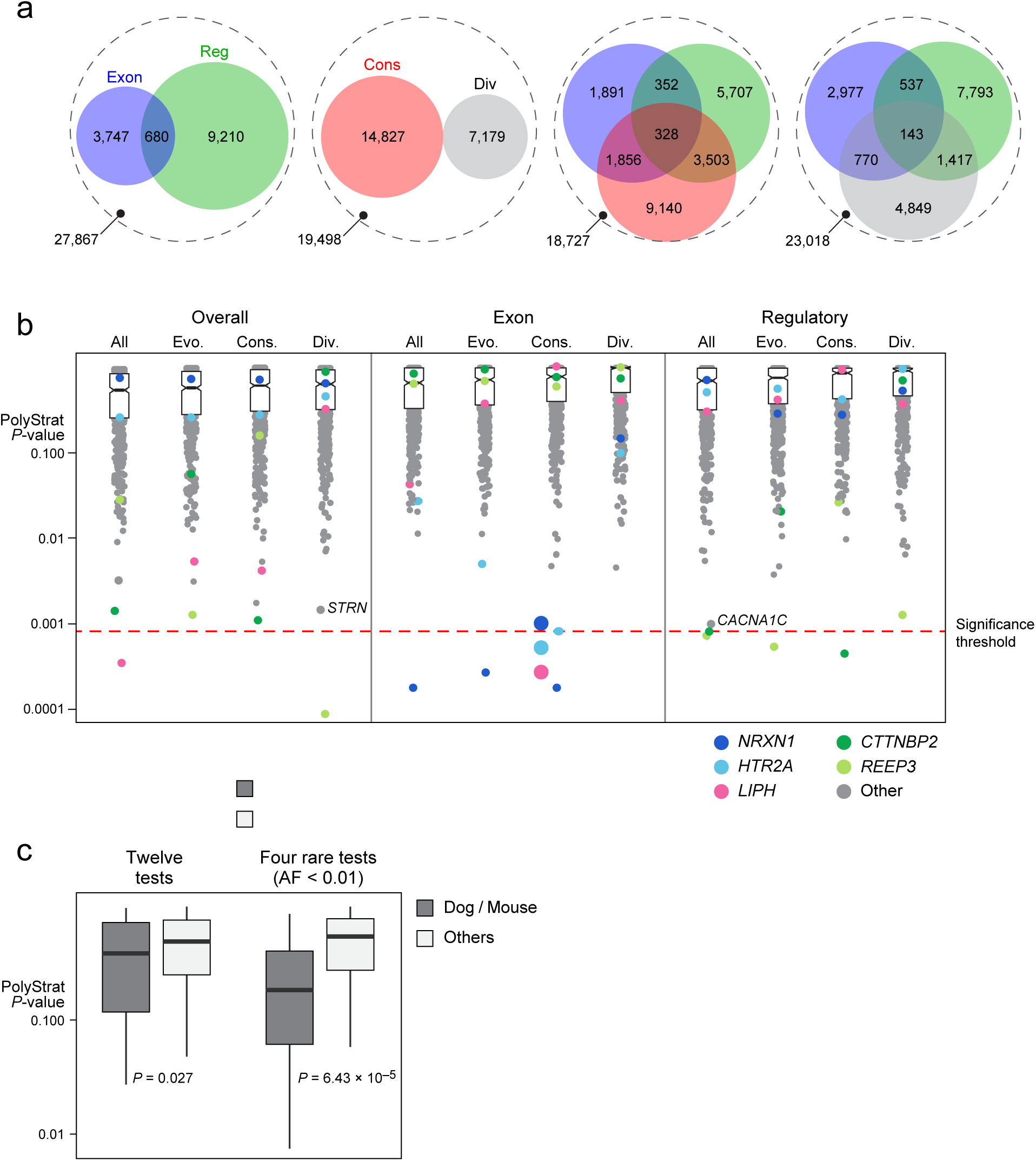
PolyStrat analysis of pooled targeted sequencing data. (a) Venn diagrams showing the number of SNPs annotated as functional and/or conserved by PolyStrat. Each of the four dashed circles represents the 41,504 total high-confidence SNPs detected. Within each circle, SNPs are stratified by their annotations. Each colorful interior circle represents SNPs annotated as exonic (blue), regulatory (green), conserved (red), or diverged (grey) bases. SNPs with multiple annotations are represented by circle overlaps, and SNPs without any of the included annotations are within the white space of the dashed outer circle. (b) PolyStrat p-values for 608 genes (circles) stratified by the 12 annotation categories tested show that just five genes (*NRXN1, HTR2A, LIPH, CTTNBP2* and *REEP3*) have p-values below the experiment-wide significance threshold after correction for multiple testing (red dashed line). Two moderately associated, OCD relevant genes discussed in the text are also noted (*STRN* and *CACNA1C*). ‘Evo.’ (=evolutionary) are SNPs either conserved (‘Cons’) or divergent (‘Div’). The vast majority of genes tested fail to exceed the significance threshold, with the median p-value for each category shown as dark black line separating two boxes representing the 25%-75% quantile. Notch in boxes shows the 95% confidence interval around median. (c) P-values for the five genes robustly implicated in animal models of OCD are significantly lower than p-values for the rest of the genes in our sequencing set (603 genes), and this difference increases when just rare variants are tested. The solid horizontal line shows median p-value, the boxed area the 25%-75% quantiles, and the vertical black lines extend from the minimum to maximum p-values observed. AF=allele frequency.

### Gene-based burden analysis

To identify genes with a significant load of non-reference alleles in OCD cases, relative to controls, we developed PolyStrat, a one-sided gene-based burden test that incorporates variant annotation. We used four variant categories: (i) all (Overall), (ii) coding (Exon), (iii) regulatory (DHS), and (iv) rare (1000 Genomes Project^53^ AF*<*0.01). Each category is further stratified by evolutionary status: (i) all; (ii) slow-evolving ‘conserved’ (Cons); (iii) fast-evolving ‘divergent’ (Div); and (iv) ‘evolutionary’ (Evo) combining (ii) and (iii). In total, PolyStrat considers 16 groups stratified by predicted function and evolutionary conservation, with more strictly defined groups likely enriched for disease-relevant variants.

PolyStrat p-values are corrected for multiple testing using a published permutation-based method that computes the empirical experiment-wide significance threshold^54,55^. This multiple testing correction method accurately measures statistical significance across correlated gene-based tests, while controlling for type 1 errors. To create an empirical null distribution, we calculated possible minimal p-values for all 9,728 tests (608 genes × 16 categories), and considered the real data that fell within the top 5% of the null distribution as significant. For most variant categories, quantile-quantile plots revealed good correspondence between observed values and the empirical null, with a small number of genes exceeding the expected distribution in a subset of the burden tests (Figures S3-S4). Our empirical correction method is more appropriate than a Bonferroni correction, which assumes that each test is independent and produces a meaningful test statistic^55^. Because the variants overlap between the categories, our tests are not independent. Furthermore, the effective number of tests run is further reduced because of an inherent limitation of the burden test, which requires sufficient variants to achieve the asymptotic properties for the test statistic^55^.

Five of the 608 sequenced genes (0.82%) show significant burdens of variants in OCD patients after correction (Table 1; Figure 1b). One, *lipase member H* (*LIPH)*, has more variants overall in OCD cases (uncorr. *p*=4 × 10^−4^, corr. p<0.026). *LIPH* encodes a protein that catalyzes the production of lysophosphatidic acid^56^. Two genes have excess coding variants: (i) *neurexin 1* (*NRXN1;* uncorr. *p*=2 × 10^−4^-3 × 10^−4^, corr. p<0.013-0.019), which encodes a synapse adhesion molecule^56^, and (ii) *serotonin receptor 2A* (*HTR2A*; uncorr. *p*=9*×*10^−4^, corr. p<0.055), an indirect target for the SSRI class of OCD medication^57^. Two genes have excess regulatory DHS variants: (i) *cortactin binding protein 2* (*CTTNBP2*; uncorr. *p*=5 × 10^−4^-9 × 10^−4^, corr. p<0.031-0.055), which modulates distribution of cortactin at the post-synapse^58^, and (ii) *receptor expression enhancing protein 3*(*REEP3)*,which may regulate cellular vesicle trafficking (uncorr. *p*=6 × 10^−4^−8 × 10^−4^, corr. p<0.036-0.049)^59^. *REEP3* is the only gene with excess divergent (potentially fast evolving) variants (uncorr. *p*=10^−4^, corr. p<0.006). We detect no genes with a significant burden of only rare variants (Figure S4).

**Table 1.**
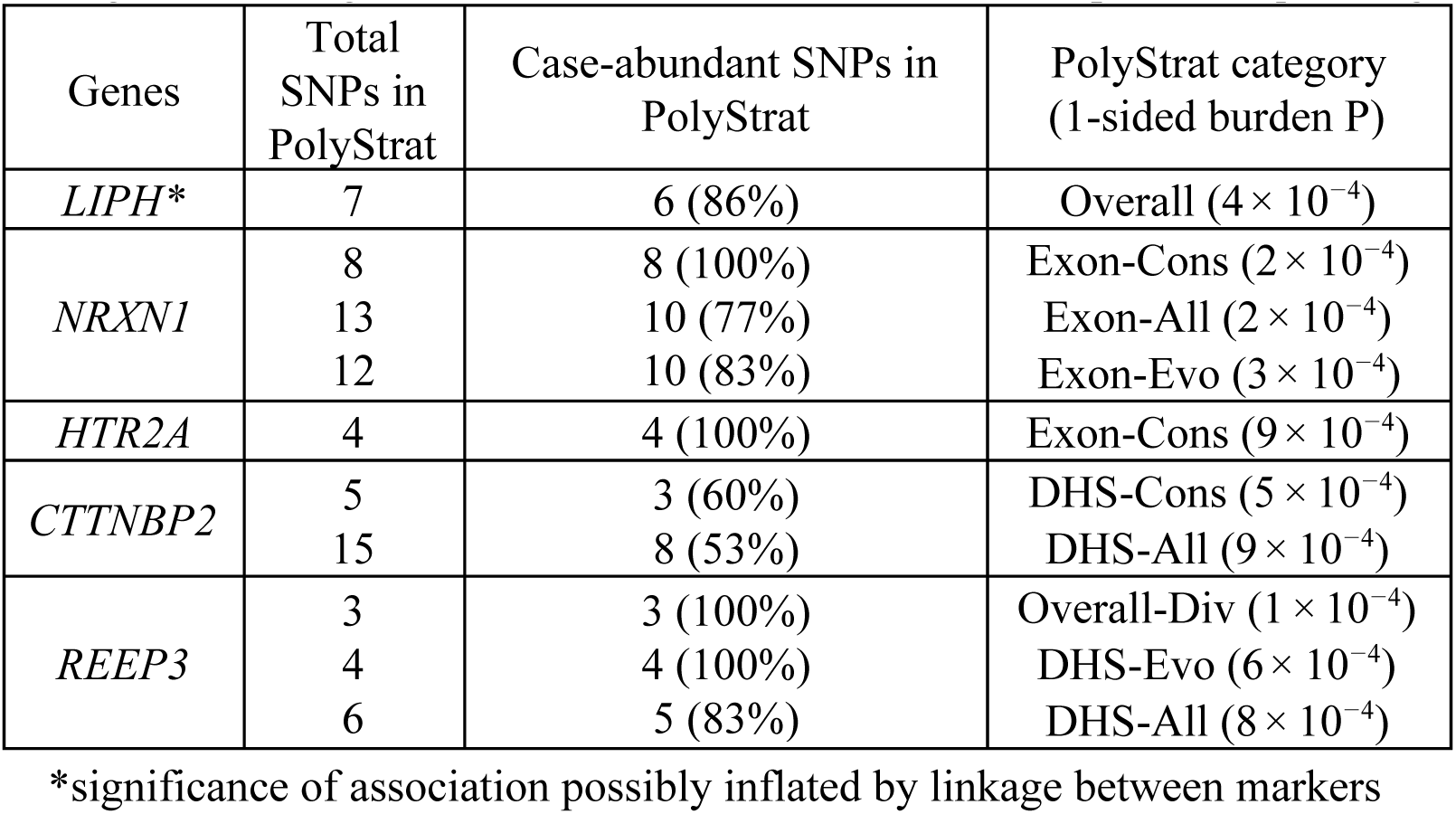
Five genes with significant variant burden in OCD cases in pooled sequencing data

| Genes | Total SNPs in PolyStrat | Case-abundant SNPs in PolyStrat | PolyStrat category (1-sided burden P) |
| --- | --- | --- | --- |
| *LIPH* * | 7 | 6 (86%) | Overall ($4 \times 10^{-4}$) |
| *NRXN1* | 8 | 8 (100%) | Exon-Cons ($2 \times 10^{-4}$) |
| | 13 | 10 (77%) | Exon-All ($2 \times 10^{-4}$) |
| | 12 | 10 (83%) | Exon-Evo ($3 \times 10^{-4}$) |
| *HTR2A* | 4 | 4 (100%) | Exon-Cons ($9 \times 10^{-4}$) |
| *CTTNBP2* | 5 | 3 (60%) | DHS-Cons ($5 \times 10^{-4}$) |
| | 15 | 8 (53%) | DHS-All ($9 \times 10^{-4}$) |
| *REEP3* | 3 | 3 (100%) | Overall-Div ($1 \times 10^{-4}$) |
| | 4 | 4 (100%) | DHS-Evo ($6 \times 10^{-4}$) |
| | 6 | 5 (83%) | DHS-All ($8 \times 10^{-4}$) |

\*significance of association possibly inflated by linkage between markers

In total, 46 SNPs contribute to the significant gene burden tests (7 in *LIPH*, 13 in *NRXN1*, 4 in *HTR2A*, 15 in *CTTNBP2*, and 7 in *REEP3*). We validated the frequencies of these SNPs in our pooled sequencing data by generating individual genotypes. After removing variants that failed Sequenom assay design or had poor call quality, we had genotype data for 37 SNPs in 571 OCD and 555 control samples (98% of the cohort) (Table S1). The allele frequencies from genotyping and sequencing were highly concordant, showing near perfect correlation (Figure S5; Pearson’s rho=0.999, *p*<2.2 × 10^−16^), validating the quality of the pooled sequencing data. For all SNPs where the direction of AF difference can be confirmed (32/37), the genotype data shifted in the same direction in cases as the sequence data. The remaining five variants cannot be confirmed either because genotypes are missing, with skew toward cases, or because an extra copy of the SNP was detected by genotyping.

We found no discernable population structure between cases and controls that would explain our significant gene associations. Allele frequencies in both case and control pools were nearly identical to 33,370 non-Finnish Europeans in ExAC (7358 SNPs, Pearson’s rho=0.995, p<2.2×10^−308^). A principal component analysis of individual genotype data for 40 ancestry informative markers found just 7 of 1152 individuals (0.6%) in the pooled sequencing cohort with potential non-European ancestry. Finally, clustering the pooled sequencing data using 1000 randomly selected SNPs, both rare and common, revealed just one potential outlier pool. To confirm that low-level ancestry differences did not affect our results, we reran PolyStrat excluding this pool and found the same five significantly associated genes.

Gene-based variant burden tests can be confounded by linkage disequilibrium (LD), whereby nearby variants tend to occur together because of population history rather than independent association with the trait of interest. Using the individual genotype data (37 SNPs), we measured LD by calculating the pairwise r^2^ for all pairs of SNPs within our five associated genes. Only one pair of SNPs, in the gene *LIPH*, was strongly linked (defined as r^2^>0.8). Thus, with the exception of the two SNPs in *LIPH*, the case-abundant variants independently contributed to the gene burden tests, and significant gene associations in *NRXN1*, *HTR2A*, *CTTNBP2* and *REEP3*were not skewed by population structure.

Under the gene-burden test, candidate genes implicated by model-organism studies (263 genes) and larger ASD studies (196 genes) were significantly more associated than genes from smaller scale human candidate gene studies (216 genes)(Kruskal-Wallis *p*=5.6×10^−15^). This is consistent with previous work showing that genes found through the smaller candidate gene studies replicate poorly^17^. It also suggests that, when a genomewide study of the disease of interest is not available, targeting genes implicated in a model organism by GWAS or by functional studies may be as effective as targeting genes implicated in a comorbid and phenotypically similar human disorder.

The five genes most strongly implicated in canine CD and murine compulsive grooming (*CDH2*, *CTNNA2*, *ATXN1, PGCP* and *Sapap3*) have significantly lower p-values than the other 603 sequenced genes (Wilcoxon unpaired, 1-sided p=2.6×10^−4^). The difference becomes more significant when only rare variants are included in the burden test (Wilcoxon unpaired, 1-sided *p*=3.2×10^−5^)(Figure 1c). This is consistent with the hypothesis that more severe disease-causing variants, while rare in humans due to negative selection, may persist at comparatively high frequencies in model organisms where selection is relaxed.

By applying the burden test across multiple genes with shared biological functions, we identified specific types of genes with a high variant load in OCD patients. We tested all 989 Gene Ontology (GO) sets that are at least weakly enriched (enrichment p<0.1) in our 608 sequenced genes (Table S2) and found two with high variant burdens: “GO:0010942 positive regulation of cell death” (uncorr. p=3 × 10^−4^, corr. p<0.03) and “GO:0031334 positive regulation of protein complex assembly” (uncorr. p=7 × 10^−4^, corr. p<0.06). Overlaying the burden test results onto the GO network topology highlights functional themes linking the enriched gene sets: regulation of protein complex assembly and cytoskeleton organization; neuronal migration; action potential; and cytoplasmic vesicle (Figure S6).

### Validation of candidate variants by genotyping

In total, our pooled sequencing identified 892 SNPs in the five genes with significant variant burdens (18 in *LIPH*, 650 in *NRXN1*, 70 in *HTR2A*, 113 in *CTTNBP2* and 41 in *REEP3*). To identify likely functional candidates, we first excluded 408 SNPs where the frequency of the non-reference, putative risk allele was higher in the controls. From the remainder, we kept SNPs that met any of the following ‘stringent’ criteria: i) single variant association p<0.05; ii) observed only in OCD cases; or iii) case frequency >2-fold higher than control frequency. We also retained SNPs that met at least two of the following ‘relaxed’ criteria: i) single variant p<0.1; ii) case frequency >1.5-fold higher than control frequency; iii) fewer than 2 observations in controls; and iv) novel (not found by the 1000 Genomes Project^53^). In total, 218 SNPs (22.3%) met our criteria for candidate variants (7 in *LIPH*, 152 in *NRXN1*, 16 in *HTR2A*, 33 in *CTTNBP2* and 10 in *REEP3*; Figure 2, Figure S7a). We ranked these SNPs by the strength of their association with OCD and selected the top 30% in each gene for further validation. This totaled 67 SNPs, including 42 rare SNPs (AF<0.01) (Figure 3a).

**Figure 2.**
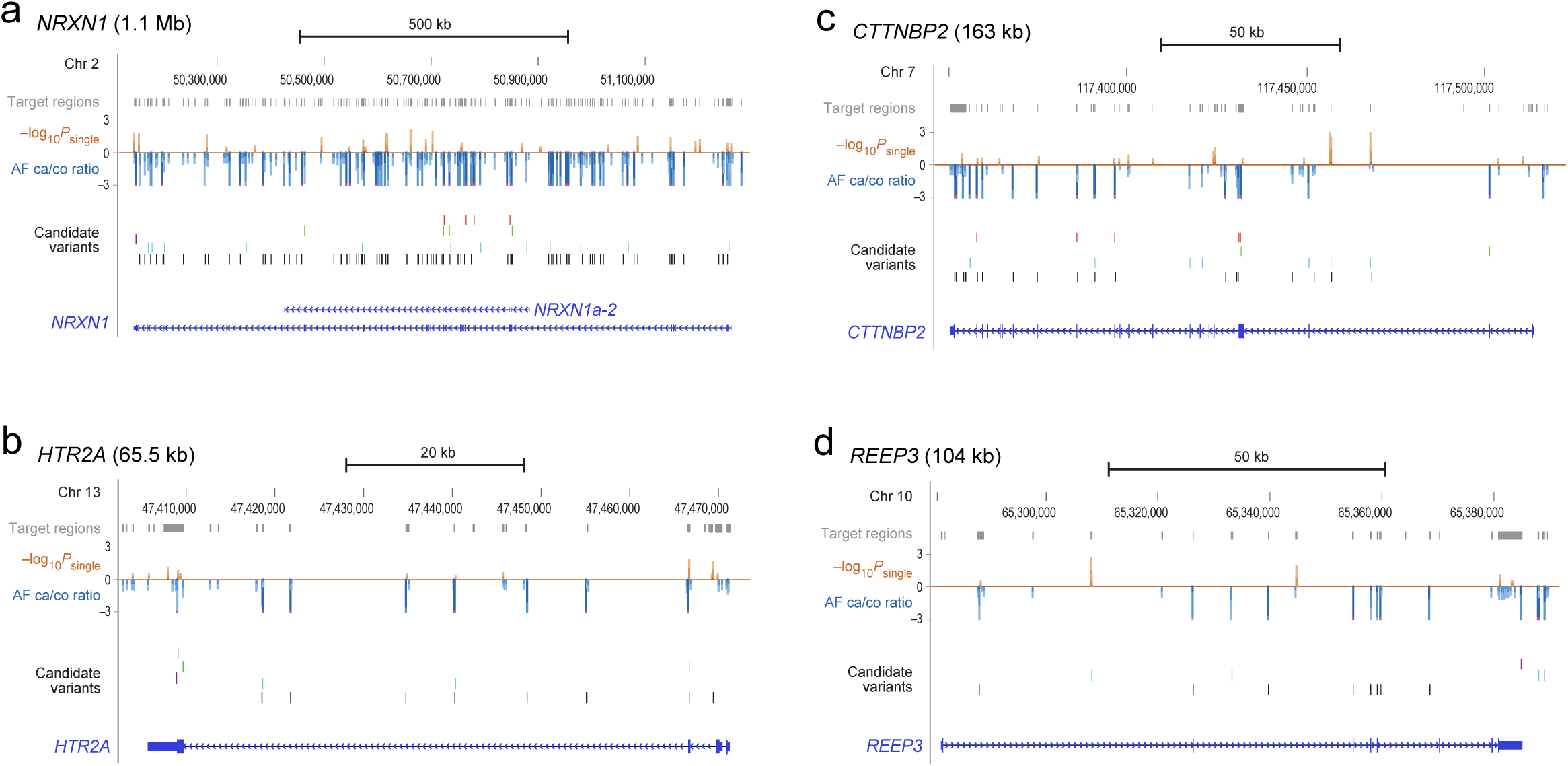
Targeted sequencing detects both coding and regulatory candidate variants in the four top-scoring genes. Sequenced “Target regions” are shown as gray boxes above the red ‘log_10_p_*single*_’ track displaying the association p-values for all detected variants and the blue ‘AF 10 *single* ca/co ratio’ track showing the ratio of OCD AF over control AF. ‘Candidate variants’ are annotated as missense (red), synonymous (green), untranslated (purple), DHS variants (blue) or unannotated (black). Lastly, the gene is shown as a horizontal blue line with exons (solid boxes) and arrows indicating direction of transcription. The highest scoring isoform of *NRXN1*, *NRXN1a-2*, is shown in (a).

**Figure 3.**
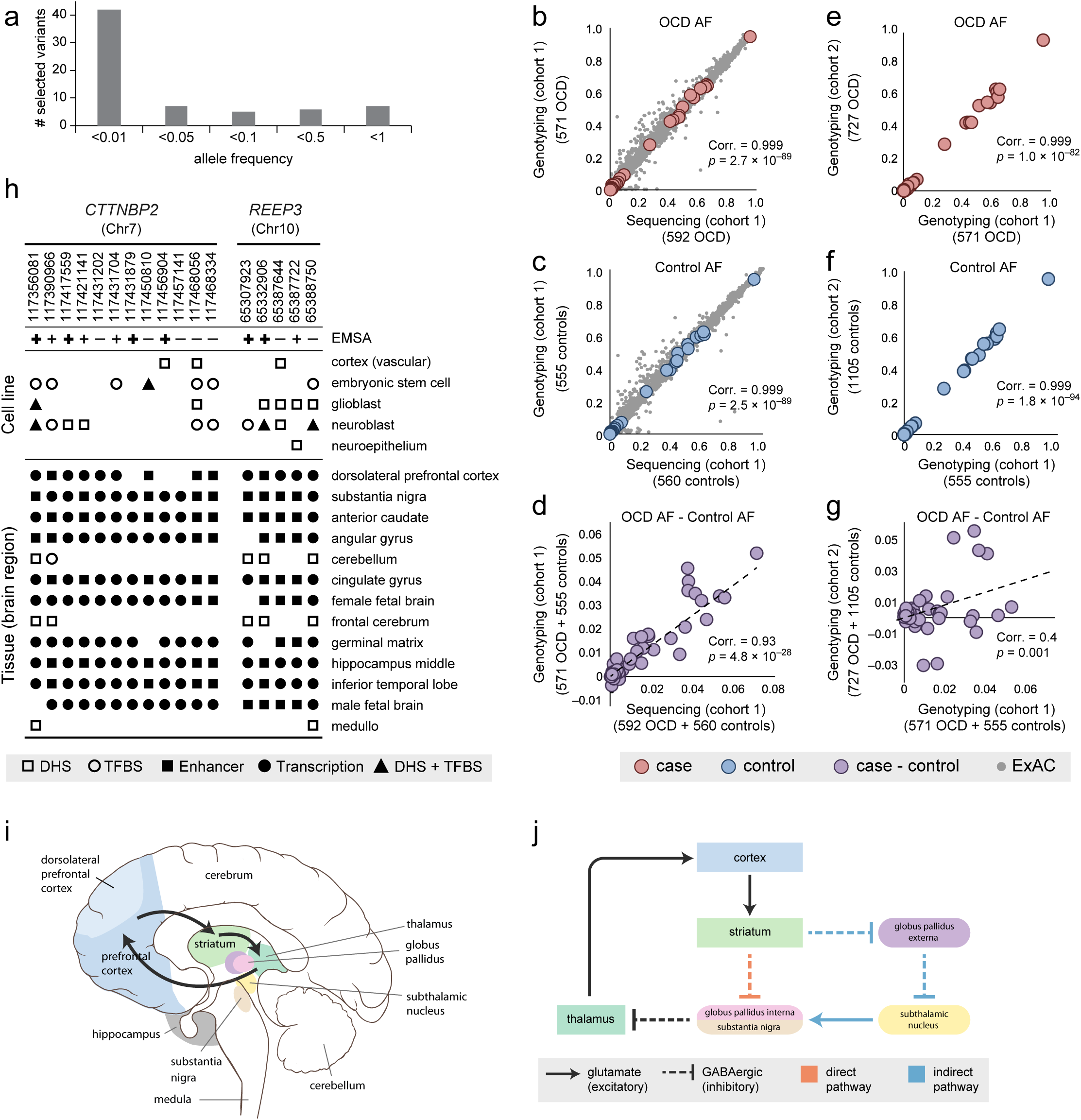
Validated top candidate variants disrupt functional elements active in brain. (a)Frequencies of the top candidate variants show that most are rare (AF<0.01) in our cohort, illustrating the value of sequencing rather than array based genotyping for detecting candidate variants. (b-g) Allele frequencies from pooled sequencing of individuals in the original cohort (cohort 1) are validated by genotyping in (b) cases and (c) controls, and by (d) allele-frequency differences between cases and controls. For the vast majority of variants, the pooled sequencing allele frequencies are also highly correlated with frequencies observed in 33,370 ExAC individuals in both cases (b, gray dots) and controls (c, grey dots). Genotyping an independent cohort (cohort 2) of cases and controls reveals genotyping allele frequencies from cohort 1 are correlated in (e) cases and (f) controls; and for (g) allele-frequency differences between cases and controls. (h) The top candidate variants in *CTTNBP2* and *REEP3*, the two genes enriched for regulatory variants, disrupt DNase hypersensitivity sites (DHS), enhancers, and transcription-factor binding sites (TFBS) annotated as functional in brain tissues and cell lines in ENCODE and Roadmap Epigenomics. All 17 variants disrupt elements active in either the dorsolateral prefrontal cortex and/or substantia nigra, which are among the (i) Brain regions involved in the CSTC circuit implicated in OCD, illustrated with black arrows. Image adapted from Creative Commons original by Patrick J. Lynch and C. Carl Jaffe, MD. (j) The CSTC circuit requires a balance between a direct, GABAergic signalling pathway and an indirect pathway that involves both GABAergic and glutamate signalling. In OCD patients, an imbalance favoring the direct over the indirect pathway disrupts the normal functioning of the CSTC circuit^61^.

We first genotyped these 67 SNPs in the same cohort used for pooled sequencing to validate the allele frequencies and get individual genotypes. After removing variants and individuals with high rates of missing data, we had 63 SNPs genotyped in 571 cases and 555 controls (98% of the cohort; genotyping rate >0.94 for all SNPs). The frequencies of these 63 SNPs are almost perfectly correlated between the pooled sequencing and genotyping data (Figure 3b, c; OCD AF, Pearson’s rho=0.999, *p*=2.7 × 10^−89^; Control AF, Pearson’s rho=0.999, *p*=2.5 × 10^−89^). The AF differences between the OCD and control cohorts are also highly concordant (Figure 3d; OCD AF - Control AF, Pearson’s rho=0.93, *p*=4.8 × 10^−28^).

We genotyped these 63 SNPs in an independent set of 727 cases and 1,105 controls of European ancestry. The OCD and control AFs in this second cohort are highly correlated with the first cohort (Figure 3e,f; OCD AF, Pearson’s rho=0.999, *p*=1.0 × 10^−82^; Control AF, Pearson’s rho=0.999, *p*=1.8 × 10^−94^). The difference in AF between cases and controls is also significantly correlated between the two cohorts (Figure 3g; OCD AF - Control AF, Pearson’s rho=0.4, *p*=0.001), and the risk allele from the first cohort is significantly more common in cases than controls in the second cohort (Wilcoxon paired 1-sided test for 63 SNPs, *p*=0.005). More specifically, 54 SNPs had a higher frequency of the non-reference allele in cases in the first cohort. Of these, 61% also had a higher frequency of the non-reference allele in cases in the second cohort. The 33 SNPs that failed to validate in either of the two cohorts had smaller allele frequency differences in the first cohort (1-sided unpaired t-test p=0.02).

In summary, the allele frequency analysis described above identified four top genes: *NRXN1*, *HTR2A*, *CTTNBP2* and *REEP3*. *LIPH* is excluded because the association is likely slightly inflated by LD and the genotyping in the second cohort did not reproduce as clearly. To validate the associations, we employed distinct strategies depending on whether the association was driven by coding variation (*NRXN1* and *HTR2A*) or regulatory variation (*CTTNBP2* and *REEP3*).

### Functional validation of regulatory variants using EMSA

For *CTTNBP2* and *REEP3*, regulatory variants give a far stronger burden test signal than does testing for either coding variants or all variants (Figure 1b). Furthermore, the three largest effect candidate variants in the combined cohort (1,298 OCD cases and 1,660 controls) alter enhancer elements in these two genes: chr7:117,358,107 in *CTTNBP2* (OR=5.2) and chr10:65,332,906 (OR=3.7) and chr10:65,287,863 (OR=3.2) in *REEP3* (Table S3). Using ENCODE and Roadmap Epigenomics data, we identified 17 candidate SNPs in *CTTNBP2* and *REEP3* likely to alter transcription factor binding sites (TFBS) and/or disrupt chromatin structure in brain and brain-related cell types^48,60^. All 17 alter enhancers or transcription associated loci active in either the substantia nigra (SN), which relays signals from the striatum to the thalamus, and/or the dorsolateral prefrontal cortex (DL-PFC), which sends signals from the cortex to the striatum/thalamus (Figure 3h,i). Both regions act in the CSTC pathway implicated by neurophysiological and genetic studies in the pathophysiology of OCD (Figure 3j)^61^.

We identified twelve candidate regulatory SNPs in *CTTNBP2* (Table 2), including: seven intronic SNPs with DHS signals in neural stem cells (SK-N-MC) or neuroblasts (SK-N-SH, BE2-C, SH-SY5Y, SK-N-SH-RA), four of which also overlap TF binding sites in the brain-derived cell lines^48^; two intronic SNPs near the top DHS variants and potentially altering the same regulatory elements; and three coding SNPs that lie within or near regulatory marks (Figure S7b). We also identified five candidate regulatory SNPs in or near *REEP3* (Table 2), including: one intronic SNP (chr10:65,307,923) that alters a DHS and GATA2 TFBS active in neuroblasts; one intronic SNP (chr10:65,332,906) that alters a DHS active in neural stem cells and GATA2, GATA3, and EP300 binding sites active in neuroblasts; three noncoding SNPs (chr10:65,387,644, chr10:65,387,722 and chr10:65,388,750) that cluster ~3kb upstream in a DHS active in multiple brain-related cells, including neuroblasts, and are seen only in OCD patients in our pooled sequencing (Figure S7c).

**Table 2.**
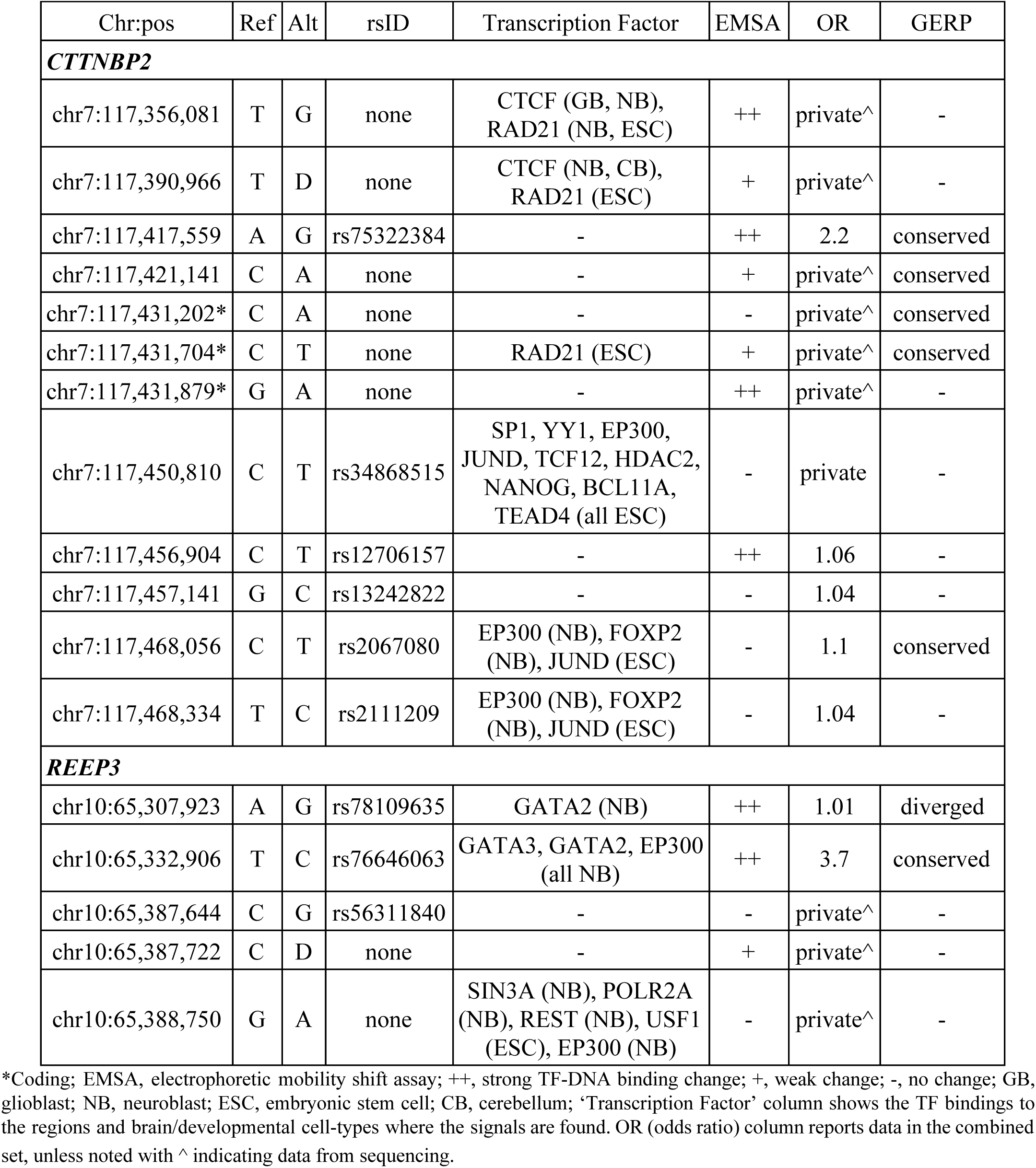
Candidate regulatory variants

We functionally tested the 17 regulatory SNPs in *REEP3* and *CTTNBP2* by introducing each into a human neuroblastoma cell line (SK-N-BE(2)) and assessing their effects on transcription-factor binding using electrophoretic mobility shift assays (EMSA). Both of the DHS SNPs in *REEP3* (chr10:65,307,923A>G and chr10:65,332,906T>C) clearly reduce specific DNA-protein binding relative to the reference allele (Figure 4a). Of the three DHS SNPs upstream of *REEP3*, only chr10:65,387,722C>del showed weak evidence of differential DNA-protein binding (Figure S8). In *CTTNBP2*, three of the seven DHS SNPs tested (chr7:117,456,904C>T, chr7:117,417,559A>G, and chr7:117,356,081T>G) clearly alter DNA-protein binding (Figure 4b), and two more, both seen only in OCD cases, (chr7:117,390,966T>del and chr7:117,421,141C>A) have weaker evidence of differential binding (Figure S8). Of the five non-DHS variants tested in *CTTNBP2*, two variants, both seen only in OCD cases, change protein binding, one strongly (chr7:117,431,879G>A; Figure 4b) and one weakly (chr7:117,431,704C>T; Figure S8).

**Figure 4.**
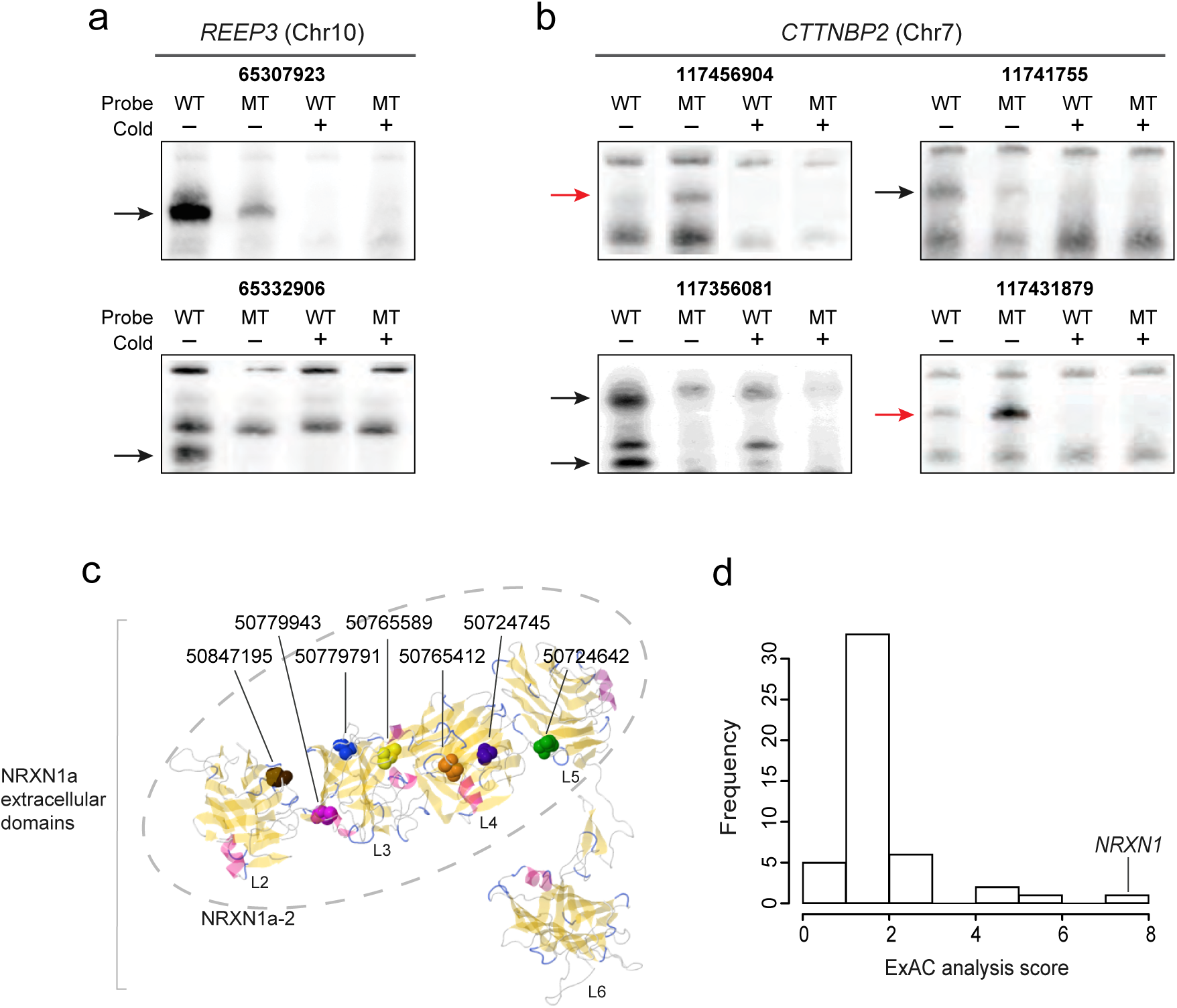
Top genes validate in functional assays, in context of known protein structure, and in comparison to ExAC. EMSA of all 17 candidate regulatory variants in (a) ***REEP3*** and (b) ***CTTNBP2***reveals that six variants either decrease (black arrows) or increase (red) protein binding when the variant sequence (MT) is compared to the reference allele (WT). The signal disappears when competing unlabeled probes (Cold +) are added, confirming the specificity of the DNA-protein binding. Contrast, size, and column orders adjusted for clear presentation. EMSA results for experimental replicates and for other candidate regulatory variants that showed weak binding changes are shown in Figure S8. (c) All seven of the candidate missense variants in ***NRXN1***, shown here as colored elements, alter the extracellular postsynaptic binding region of our top-scoring protein isoform NRXN1a-2^77,103,104^. Of the six extracellular LNS (laminin, nectin, sex-hormone binding globulin) domains in the longer isoform NRXN1a, five with known protein structure are shown and labeled L2-L6. NRXN1a-2 includes four of these domains (L2-L5; dashed ellipse). (d) The isoform-based test comparing OCD cases to ExAC finds ***NRXN1*** as the top scoring gene with genomewide significance. The ExAC analysis score is defined as the ratio of χ^2^ statistics 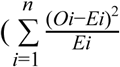 where ***n*** = total number of isoforms, ***Oi*** = number of non-reference alleles observed in isoform ***i***, ***Ei*** = number of non-reference alleles expected from ExAC in isoform ***i***) between OCD vs. ExAC comparison and control vs. ExAC comparison.

The high rate of functional validation by EMSA demonstrates that screening using both regulatory and evolutionary information is remarkably effective in identifying strong candidate OCD risk variants. In total, eight of 12 tested DHS SNPs (67%) show evidence of altered protein binding relative to the reference allele, despite testing them within a single cell line at a single time point under basic binding conditions (Table 2). This includes two SNPs with high ORs in the full genotyping datasets that strongly disrupt specific DNA-protein binding (Chr10:65,332,906 with OR=3.7; chr7:117,417,559 with OR=2.2). Two of five non-DHS SNPs (40%) also show altered binding, illustrating that DHS mark alone is a powerful but imperfect predictor of regulatory function. Both of these SNPs alter highly constrained elements (SiPhy score 8.7), while only one of the three non-DHS SNPs is constrained. This suggests that incorporating both DHS and conservation identifies functional regulatory variants with more specificity, consistent with other published results^62^.

### Validation of coding variants using ExAC

In contrast to the regulatory variant burden found in *CTTNBP2* and *REEP3, NRXN1* and *HTR2A* showed significant PolyStrat signals when only coding variants were considered, suggesting a role for protein-coding changes in these genes in OCD. Among the coding SNPs detected in *NRXN1*, twelve met our candidate SNP selection criteria. Of the twelve, seven are missense (Table 3), of which three are novel SNPs and private to OCD cases in our data^49^. The remaining four missense SNPs are known rare SNPs with OCD AFs ranging 0.002-0.009. All these seven SNPs change amino acids in laminin G or EGF-like domains that are important in binding to other synaptic adhesion molecules at the post-synapse, hence potentially affecting the involvement of NRXN1 in synapse formation and maintenance (Figure 4c).

**Table 3.** Coding variants in NRXN1 and HTR2A

| Chr:pos | Ref | Alt | rsID | OCD allele freq. | Ctrl allele freq. | ExAC allele freq. | Amino acid change | Candidate variant from sequencing |
| --- | --- | --- | --- | --- | --- | --- | --- | --- |
| ***NRXN1*** |  |  |  |  |  |  |  |  |
| chr2:50,149,133 | C | T | rs113380721 | 0.0019 | 0.0030 | 0.0033 | Syn | no |
| chr2:50,149,214 | A | G | rs112536447 | 0.0001 | 0.0010 | 0.0003 | Syn | no |
| chr2:50,280,604 | T | C | rs79970751 | 0.0088 | 0.0070 | 0.0058 | Syn | no |
| chr2:50,463,984 | G | A | rs147580960 | 0.0009 | 0.0006 | 0 | Syn | yes |
| chr2:50,464,065 | C | T | rs80094872 | 0.0009 | 0.0003 | 0* | Syn | yes |
| chr2:50,699,479 | G | A | rs75275592 | 0.0009 | 0.0013 | 0.0010 | Syn | no |
| chr2:50,723,068 | G | A | rs56402642 | 0.0034 | 0.0001 | 0.0018 | Syn | yes |
| chr2:50,724,642 | A | G | none | 0.0009 | 0 | 0 | I>T | yes |
| chr2:50,724,745 | G | T | rs201818223 | 0.0017 | 0.0009 | 0.0011 | L>M | yes |
| chr2:50,733,745 | G | C | rs147984237 | 0.0017 | 0 | $1.52 \times 10^{-5}$ | Syn | yes |
| chr2:50,765,412 | G | T | rs56086732 | 0.0089 | 0.0028 | 0.0056 | L>I | yes |
| chr2:50,765,589 | T | C | rs200074974 | 0.0016 | 0 | 0.0011 | I>V | yes |
| chr2:50,779,791 | C | T | none | 0.0008 | 0 | 0* | A>T | yes |
| chr2:50,779,943 | T | C | none | 0.0009 | 0 | 0* | N>S | yes |
| chr2:50,847,195 | G | A | rs78540316 | 0.0077 | 0.0036 | 0.0043 | P>S | yes |
| chr2:50,850,686 | G | A | rs2303298 | 0.0107 | 0.0013 | 0.0038 | Syn | yes |
| ***HTR2A*** |  |  |  |  |  |  |  |  |
| chr13:47,409,048 | G | A | rs6308 | 0.0036 | 0.0012 | 0.0023 | A>V | yes |
| chr13:47,409,701 | G | A | rs141413930 | 0.0020 | 0.0012 | 0.0026* | Syn | yes |
| chr13:47,409,149 | T | A | rs35224115 | 0.0019 | 0.0047 | 0.0044 | Syn | no |
| chr13:47,466,622 | G | A | rs6305 | 0.0386 | 0.0225 | 0.0266** | Syn | yes |

ExAC allele freq. shown for NFE population
\*Excluded from ExAC analysis because low-confidence call from pooled sequencing
\*\*Excluded from ExAC analysis because frequency > 0.01

Two of the three *HTR2A* coding SNPs, a missense and a synonymous SNP, are located in the last coding exon (205 to 471aa in the UniProt protein P28223). The missense SNP (rs6308) is located in the cytoplasmic domain, which contains the PDZ-binding motif where protein interacting partners bind. Missense variants in this region have been shown to alter protein binding properties and disrupt the interaction of PDZ-based scaffolding proteins at the synapse, suggesting that the *HTR2A* coding variants we find may affect binding affinity or specificity^63^.

We next sought to improve our statistical power by combining our pooled sequencing data with the publicly available genomic data from ExAC^33^. The associations of *CTTNBP2, REEP3, NRXN1*, and *HTR2A* with OCD are experiment-wide significant, but do not reach the genomewide significance threshold of p<2.5 × 10^−6^ (~20,000 human genes)^55^. When discovery and validation cohorts are combined, our strongest association, to *NRXN1*, is *p*=5.1 ×10^−5^ (Fisher’s combined *p*). For genes with a burden of coding variants (*NRXN1* and *HTR2A)*, ExAC offers an opportunity to assess the significance of the variant burden in OCD cases compared to a cohort of 33,370 non-Finnish Europeans. We note that such a comparison was not possible for *CTTNBP2* and *REEP3*, for which the associated variants are predominantly noncoding and thus not present in ExAC.

The public ExAC database only provides allele frequencies for variants; our pooled sequencing data has the same constraint. Thus, permutation based approaches (e.g. case/control label swapping) that require individual-level genetic data are not possible. Instead, we used our own controls to find a reliable null model. Because our allele frequencies are almost perfectly correlated with ExAC (Pearson’s rho=0.995, p<2.2 × 10^−308^ for 7358 shared variants; Figure 3b,c), we expect no association when comparing our controls to ExAC. However, using the Fisher’s Exact test gives highly significant association signals even for this “null” comparison, due to the extremely large size of the ExAC cohort^64^.

We instead use an isoform-based test that compares the distribution of variants across different isoforms of the same gene. By incorporating a within-gene comparison for assessing significance, this effectively controls inflation in the null case. We first calculate, using ExAC, the number of rare (AF<0.01), non-reference, coding alleles within each isoform. We then used the χ^2^ goodness-of-fit test to compare the distribution of variants across isoforms in our data and ExAC. Of 542 genes with more than one isoform, we saw no significant difference between our control data and ExAC for over 90% (493 genes had corr. p>0.05). Focusing on the subset of 66 genes with rare allele counts that could produce nominally significant PolyStrat scores, *NRXN1* had the largest difference between cases and ExAC (χ^2^=82.3, df=16, uncorr. p=6.37 × 10^−11^; corr. p=1.27 × 10^−6^) and no difference between controls and ExAC (χ^2^=10.5, df=16, uncorr. p=0.84)(Figure 4d). We note that *HTR2A*, the other gene enriched for coding variants, had only two SNPs in cases, which was too little information to use the isoform test.

The significant association of *NRXN1* reflects an exceptional burden of variants in one of the 17 Ensembl isoforms defined for the gene. *NRXN1a-2*, which contains all 12 candidate coding SNPs, had the largest deviation between observed and expected variant counts, with a residual at least 1.4x higher than any other isoform (*NRXN1a-2*=22.3, *NRXN1-001*=16.3; median=5.15). After adjusting for the residuals from the ‘null’ control data and ExAC comparison, the *NRXN1a-2* residual is still 1.3x higher (OCD residual/Control residual *NRXN1a-2*=5.34, *NRXN1-014*=4.04).

## DISCUSSION

By analyzing sequencing data for genes previously implicated in OCD, and then prioritizing variants according to their functional and conservation annotations, we identify four genes with a reproducible variant burden in OCD cases. Two genes, *NRXN1* and *HTR2A* (Table 3), have a burden of coding variants, and the other two, *CTTNBP2* and *REEP3* (Table 2), have a burden of conserved regulatory variants. Notably, all four genes act in neural pathways linked to OCD, including serotonin and glutamate signaling, synaptic connectivity, and the CSTC circuit^65,66^, offering new insight into the biological basis of compulsive behavior (Figure 5).

**Figure 5.**
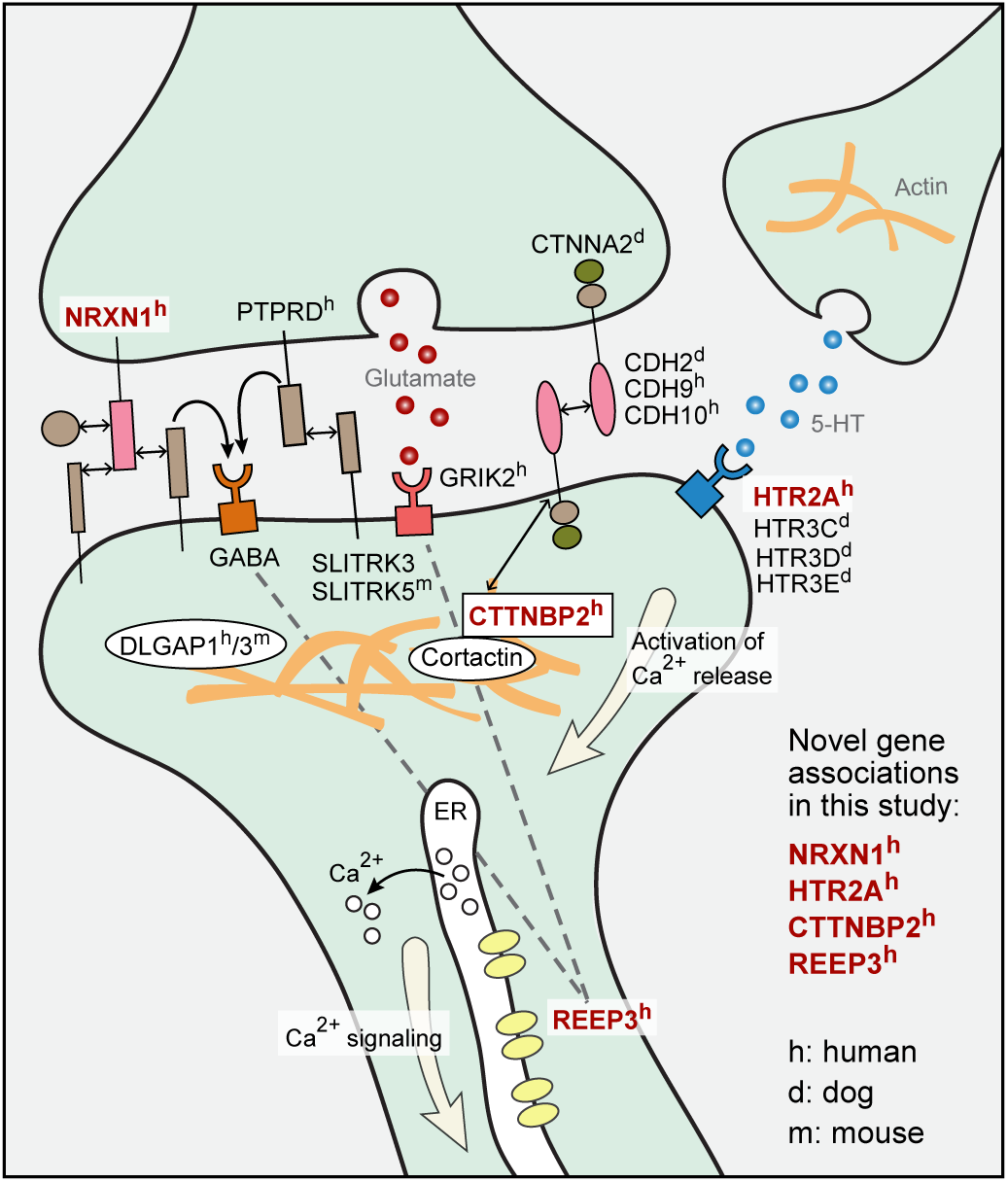
All four top candidate genes function at the synapse and interact with proteinsimplicated in OCD by previous studies in human (superscript h), dog (superscript d) and mouse (superscript m)^65,66^. Genes identified in this study are shown in red. Solid lines indicate direct interactions, and dashed lines indicate indirect interactions.

We use three independent approaches to validate our findings: (1) For the top candidate SNPs, allele-frequency differences from targeted sequencing data are confirmed by genotyping of both the original cohort (Figure 3d) and a larger, independent cohort (Figure 3g). (2) For the two genes with a burden of coding variants (*NRXN1* and *HTR2A)*, comparison of our data to 33,370 population-matched controls from ExAC^33^ reveals genomewide-significant association of *NRXN1* with OCD (χ^2^=82.3, df=16, uncorr. p=6.37 × 10^−11^; corr. p=1.27 × 10^−6^). (3) For the two genes with burden of regulatory variants (*REEP3* and *CTTNBP2)*, we show that at least one third of candidate SNPs are functional, altering protein to DNA binding in a neuroblastoma cell line (Figure 4a).

Comparison of our approach to existing methods illustrates its unique advantages, and offers a deeper understanding of how its two key features — targeted sequencing, and incorporation of functional and conservation metrics — permit identification of significantly associated genes using a cohort smaller than those that have previously failed to yield significant results.

Our targeted-sequencing strategy permits identification of both coding and regulatory variants, and both common and rare variants, at only a fraction of the cost of whole-genome sequencing (WGS). For the modest-size cohort in this study, WGS would cost ~$2.5M at current prices, 40-fold more than our pooled-sequencing approach. Even without pooling, our targeted sequencing would cost 4-fold less than WGS. Whole-exome sequencing would cost approximately the same as targeted sequencing, but would miss the regulatory variants which explain most polygenic trait heritability^31,32^. By using existing information to prioritize a large set of genes likely enriched with genes involved in OCD, and then performing targeted sequencing of functional elements, our approach enhances causal-variant detection and thus statistical power, although at the cost of missing any OCD-associated genes not included as candidates as well as potential distant regulatory elements.

With targeted sequencing data, we can detect associations to rare variants. This feature is especially critical for study of diseases that, like OCD, may reduce fitness, as negative selection drives down the frequency of deleterious variants that are passed on to the next generation ^67^. Genotype array data sets, and even imputed data sets, are not as effective, as they are missing many rare variants. In our data set, 80% of variants driving significant associations have allele frequencies < 0.05; one of the densest whole genome genotyping arrays currently available, the Illumina Infinium Omni5 with 4.3M markers, contains only half of these variants(Table S1)^2,3^. In addition, 60% of our variants have allele frequencies < 0.01, and would be missed even through imputation with 1000 Genomes and UK10K^53,68,69^.

To analyze the targeted sequencing data, we developed PolyStrat, a new analytical method that gains statistical power by leveraging public evolutionary and regulatory data for the human genome. Given sequence data from cases and controls, PolyStrat starts by filtering out variants that are less likely to be functional, and then performs a gene burden test with the remaining variants. In contrast to other gene-based approaches, which focus on ultra-rare, protein-damaging variants, PolyStrat considers variants with diverse allele frequencies. By considering a large subset of variants enriching for functional changes, PolyStrat gains statistical power for identifying genes with an excess of variants in cases, within a standard gene-level burden framework.

PolyStrat is particularly advantageous for identifying associated genes in studies with smaller cohorts. It tests for association at the gene level, and thus requires statistical correction only for the ~20,000 genes in the genome, rather than testing individual variants, which requires correcting for millions of independent variants in the genome. PolyStrat further increases power by optimizing the key parameters in a gene-based test, namely increasing the proportion of causal variants in each gene, and including variants with higher allele frequencies and/or larger effect sizes^70^. PolyStrat’s functional annotation includes ~82 times more functional variants than an approach focusing on protein-damaging variants (27,626 by PolyStart vs. 335 by PolyPhen2^71^ in our data). PolyStrat increases the proportion of functional variants first by targeting only the ~10% of sequence in each gene that is coding or evolutionarily constrained, and subsequently removing another 33% of variants unlikely to be functional.

We observed that the frequencies and effect sizes of the candidate causal variants detected by PolyStrat are consistent with what we expect from simulations. Simulation suggests that with 200-700 cases, we should have 90% power to detect associated genes with combined allele frequencies and effect sizes similar to our four genes^72^. The candidate causal variants in *NRXN1* have combined frequency of 0.022 and OR of 2.4 and would be detectable in ~600 cases with 90% power; variants in *HTR2A* (combined frequency=0.03, OR=1.56) would be detectable with ~700 cases; rare variants in *CTTNBP2* (combined frequency=0.003, OR=4.7) would be detectable with ~500 cases; and variants in *REEP3* (combined frequency=0.04, OR=2.11) would be detectable with ~200 cases.

Previous research on the four genes identified by PolyStrat suggests they act in brain function relevant pathways, and harbor variants that could potentially alter OCD risk (Table 4)^65,66^:

**Table 4.** Summary of top genes

|  | *NRXN1* | *REEP3* | *CTTNBP2* | *HTR2A* |
| --- | --- | --- | --- | --- |
| **Gene Product/**<br>**Brain**<br>**Relevance** | synaptic cell-adhesion protein/ synapse functioning and development in cortical-striatal pathway<sup>73,74</sup> | microtubule-binding protein/ possible role in synaptic plasticity, calcium signaling, shaping tubular ER membranes in neurons<sup>78</sup> | cortical actin (cortactin) binding protein/ synaptic maintenance<sup>81</sup> | G-protein coupled serotonin receptor/ cortical neuron excitation<sup>101</sup> |
| **Disease**<br>**Relevance** | neurodevelopmental disorders incl. ASD<sup>75,76</sup> | ASD<sup>59</sup> | interacts with CDH2, implicated in canine CD<sup>10,11,82</sup> | ASD, OCD<sup>83</sup>, canine CD (5-HT3 receptors)<sup>84</sup> |
| **Reason for**<br>**Inclusion**<br>**as Candidate\*** | 1) model (mouse) organism gene;<br>2) ASD gene | 2) ASD gene | 1) model (mouse) organism gene | 1) model (dog) organism gene;<br>3) human candidate - SSRI target |
| **Type of Burden**<br>**Identified** | coding variants - missense variants exclusive to one isoform, *NRXN1a-2* | regulatory variants | regulatory variants | coding variants - missense variant in perfect linkage to a common variant rs6314 associated with response to SSRIs<sup>57,85</sup> |
| **Validation of**<br>**Variants**<br>**Identified**<br>**in Present**<br>**Study** | by comparison to ExAC - genome-wide significant association | by EMSA - disrupt regulatory elements bound by various TF, including GATA2 | by EMSA - alter epigenetic marks active in the cortico-striatal pathway | by genotyping independent cohort; too few polymorphic sites for validation with ExAC |
| **Hypothesized**<br>**Impact of**<br>**Variants**<br>**Identified** | inaccurate cellular localization of NRXN1, or altered binding competition to its partner, modifying synaptic adhesion<sup>77</sup> | *REEP3* expression in GABA neurons inhibited by variants that reduce GATA2 binding<sup>79</sup>, leading to excitatory/inhibitory imbalance in CSTC circuit<sup>61,80</sup> | altered *CTTNBP2* expression in cortico-striatal circuit in brain<sup>102</sup> | altered binding affinity of HTR2A, changing the activation of downstream calcium signaling in neurons<sup>63</sup> |

\*For category types, see Results

***NRXN1***. *NRXN1* encodes the synapse cell adhesion protein neurexin 1α, which acts in the MSN microcircuit in the cortico-striatal neural pathway and is critical for synapse development^73,74^. Mutations in this gene are causally linked to ASD and other psychiatric diseases, with individual *NRXN1* isoforms implicated in distinct neuropsychiatric disorders^75,76^(Figure 5). NRXN1 is functionally similar to cadherins that mediate synaptic adhesion, including the canine CD gene *CDH2* and *CDH9/CDH10*, top human OCD candidates ^3,10,11^ (Figure 5). Our candidate variants in the top-scoring isoform NRXN1a-2 have potential to exert a dominant negative effect by inaccurate cellular localization or by competing for neurexin 1α interacting partners, particularly through neurexophilins, potentially leading to the synaptic dysfunction observed in OCD^77^.

***REEP3***. Rare mutations in the synaptic plasticity and calcium signaling gene *REEP3* are implicated in autism^59^. REEP3 proteins help shape the tubular ER membranes that extend throughout highly polarized cells such as neurons^78^. We functionally validated 2 out 5 variants tested by EMSA. Both validated variants disrupt regulatory elements active in the cortico-striatal neural pathway that are bound by multiple TFs, including GATA2 (Table 2, Figure 3h, 4a). Gata2 is required to actuate GABAergic neuron-specific gene expression, rather than a glutamatergic phenotype, in the mouse embryonic midbrain^79^. Both inhibitory GABA and excitatory glutamate neurons are critical components of the CSTC circuit(figure 3j)^61,80^. We hypothesize that, by inhibiting expression of ***REEP3*** in GABA neurons, variants disrupting GATA2 binding could cause an imbalance between excitatory and inhibitory neurons in the CSTC circuit.

***CTTNBP2.*** *CTTNBP2*, which had a burden of regulatory variants in our cases, was included in the set of targeted genes because of its role in regulating postsynaptic excitatory synapse formation and maintenance in the cortico-striatal brain circuit^81^ (Figure 5). CTTNBP2 interacts with proteins encoded by striatin (*STRN*), which had a near-significant burden of variants in this study (uncorr. p=0.0016, corr. p<0.1; Figure 1b) and *CDH2*, which is implicated in canine CD^10, 11, 82^. We used EMSA to test 12 variants in *CTTNBP2*, and show four of the variants clearly disrupt protein binding (Table 2). All four alter epigenetic marks active in the key structures of the cortico-striatal neural pathway (Table 2, Figure 3h and 4a). Regulatory variants in *CTTNBP2* could affect OCD risk by altering the expression of this critical gene in the cortico-striatal circuit of the brain.

***HTR2A.*** *HTR2A*, implicated in ASD and in candidate gene studies of OCD, encodes a G-protein coupled receptor, serotonin (5-HT) receptor 2A, that is expressed widely throughout the central nervous system, including the prefrontal cortex. In humans, meta-analysis of 19 datasets for *HTR2A* indicated a robust association of *HTR2A* with OCD^83^, and in dogs a related serotonin receptor cluster (*HTR3C/HTR3D/HTR3E*) is associated with severe canine CD ^84^(Figure 5). We found three coding variants that could potentially alter the binding affinity of HTR2A (Table 3)^63^. Most notably, the rare missense candidate variant rs6308 (AF=0.004 in 1000G CEU population) is perfectly linked (D’=1) to a common variant (rs6314) associated with response to SSRIs ^57,85^.

Taken together, our top four associated genes, and our pathway analysis, implicate particular neuronal functions in OCD:

**Cell-adhesion**. Synaptic cell adhesion molecules help establish and maintain contact between the presynaptic and postsynaptic membrane, and are critical for synapse development and neural plasticity. *NRXN1* encodes a cell adhesion molecule predominantly expressed in the brain, and *CTTNBP2* regulates cortactin, another such molecule. Our pathway analysis showed that genes involved in ‘regulation of protein complex assembly’ and ‘cytoskeleton organization’ were enriched for variants in OCD patients. These results echo earlier findings of relevance to compulsive disorders for cell-adhesion genes in dogs (*CDH2* and *CTNNA2*), mice (*Slitrk5)*, and in humans (*DLGAP1, PTPRD* and *CDH9/CDH10*)^2,3,10,86^(Figure 5).

**Excitatory and inhibitory signaling**. An imbalance of excitatory glutamate and inhibitory GABAergic neuron differentiation underlying OCD has been proposed in the CSTC-OCD model (Figure 3j)^61^. *NRXN1* induces differentiation of GABA/glutamate postsynaptic specialization^87^. *REEP3* might be involved in GABA/glutamate neuron-specific expression (Table 4)^88^. A top gene from OCD GWAS, *PTPRD*, promotes glutamatergic synapse differentiation through this pathway^3^. Moreover, we find an overall burden of variants in genes regulating cell death and apoptosis (Table S2) and in telencephalic tangential migration (TTM), a major neuronal migration event during cerebral development involving extensive cell death. TTM forms neuronal connections between the key structures of CSTC circuit, and if perturbed could result in an imbalance of excitatory-inhibitory signals and abnormal connectivity within the CSTC circuit^89^.

**Serotonin signaling**. SSRIs are the most effective treatment available for OCD, suggesting serotonergic pathways are involved in the disease. *HTR2A* encodes a serotonin receptor, and allelic variation in *HTR2A* alters response to SSRIs (Table 4)^57,90,91^. In addition, both *REEP3* and *CACNA1C*, a high scoring gene in this study (Figure 1) also significantly associated with schizophrenia, act in calcium signalling, a downstream pathway of *HTR2A*^92–94^. A meta-analysis of >100 OCD genetic association studies found strong association to both *HTR2A* and the serotonin transporter gene *SLC6A4*^83^. In dogs, a serotonin-receptor locus is associated with severe CD^84^.

Our findings also suggest broad principles that could help to guide studies of other polygenic diseases. We find that genes associated in selectively bred model organisms are more likely to contain rare, highly penetrant variants. The five genes we found to be most strongly associated with compulsive behaviors in dog and mouse (*CDH2*, *CTNNA2*, *ATXN1, PGCP* and *Sapap3)* were significantly more enriched for rare variants in human patients than the other 603 genes we targeted, although they did not individually achieve significance (Figure 1c). We propose that the enrichment of rare variants in humans reflects natural selective forces limiting the prevalence of severe disease-causing variants. Such forces are less powerful in selectively bred animal populations. Because risk variants identified through animal models are anticipated to be rare in humans, replication will require either family-based studies, or cohorts of magnitude not currently available.

We also find that the ratio of coding to regulatory variants is positively correlated with the gene’s developmental importance. While single-gene p-values from PolyStrat tests are positively correlated across variant categories, as is expected given the overlaps between different variant categories (Figure 1a, Figure S9), this pattern breaks down for our four significantly associated genes. *NRXN1* and *HTR2A*, which have burdens of coding variants, score poorly on regulatory-variant tests; *CTTNBP2* and *REEP3*, which have burdens of regulatory variants, score poorly in coding-variant tests (Figure 1b). This is consistent with the ExAC study showing that genes critical for viability or development do not tolerate major coding changes^33^. In that study, the authors estimate that it is highly probable that *CTTNBP2* and *REEP3* would be intolerant of homozygous loss of function variants (pRec = 0.99999015 and pRec = 0.953842585, respectively), while *HTR2A* (pRec=0.225555783) and, most notably, *NRXN1*(pRec=5.13x10^-5^) would be far more tolerant. The enrichment for regulatory variants in *CTTNBP2* and *REEP3* that we find suggests that these genes may be able to tolerate variants with more subtle functional impacts, such as expression differences in specific cell types or developmental stages.

### Outlook

With technological advances in high-throughput sequencing, there is an increasing focus on identifying causal genetic variants that underlie diseases as a first step toward targeted therapies^95^. However, existing approaches have notable limitations. Whole genome sequencing is prohibitively expensive on large cohorts, while cost-saving whole-exome sequencing does not capture the regulatory variants underlying complex diseases^32^. Leveraging existing genomic resources can increase power to find causal variants through meta-analysis and imputation, but these resources are heavily biased towards a few populations^96^. Without new approaches, advances in precision medicine will predominantly benefit those of European descent^97^.

Here, we describe an approach that uses prior research, targeted sequencing and a new analysis method to efficiently identify genes and individual variants associated with complex disease risk in any population. In a modest-size cohort of OCD cases and controls we find associations driven by both coding and regulatory variants, highlighting new potential new therapeutic targets. Our method holds promise for elucidating the biological basis of complex disease, and for extending the power of precision medicine to previously excluded populations.

## METHODS

### Study design

We designed and carried out the study in two phases. In the first, discovery phase, we performed targeted sequencing of 592 individuals with DSM-IV OCD^98^ and 560 controls of European ancestry, and tested association for OCD at single variant-, gene-, and pathway-level. In the second, validation phase, we employed three distinct analyses. (1) We genotyped both the original cohort and a second, independent cohort containing 1834 DNA samples (729 DSM-IV OCD cases and 1105 controls) of European ancestry, including a total of 2986 individuals (1321 OCD cases and 1665 controls) to confirm the observed allele frequencies in the discovery phase. (2) We compared our sequencing data with 33,370 population-matched controls from ExAC to confirm the gene-based burden of coding variants as well as allele frequencies. (3) We performed EMSA to test whether our candidate variants have regulatory function.

### Targeted regions

We targeted 82,723 evolutionarily constrained regions in and around 608 genes, which included all regions within 1kb of the start and end of each of 608 targeted genes with SiPhy evolutionarily constraint score >7, as well as all exons^46^. For the intergenic regions up and downstream of each gene, we used constraint score thresholds that became more stringent with distance from the gene.

### Pooled sequencing and variant annotation

Groups of 16 individuals were pooled together into 37 case pools and 35 control pools and barcoded. Targeted genomic regions were captured using a custom NimbleGen hybrid capture array and sequenced by Illumina GAII or Illumina HiSeq2000. Sequencing reads were aligned and processed by Picard analysis pipeline (http://broadinstitute.github.io/picard/). Variants and AFs were called using Syzygy^25^ and SNVer^47^. We used ANNOVAR^50^ to annotate variants for RefSeq genes (hg19), GERP scores, ENCODE DHS cluster, and 1KG data.

### Genotyping

SNP genotyping was performed using the Sequenom MassARRAY iPLEX platform and the resulting data was analyzed using PLINK1.9^99^.

### EMSA

For each allele of the tested variants, pairs of 5’-biotinylated oligonucleotides were obtained from IDT Inc. (Table S4). Equal volumes of forward and reverse oligonucleotides (1 pmol/ul) were mixed and heated at 95°C for 5 minutes and then cooled to room temperature. Annealed probes were incubated at room temperature for 30 minutes with SK-N-BE(2) nuclear extract (Active Motif). The remaining steps followed the LightShift Chemiluminescent EMSA Kit protocol (Thermo Scientific).

### Statistical analysis

For gene-/pathway-association, we employed a simple burden test approach, which has been used before^54^. Briefly, we used the sum of the differences of non-reference allele rates between cases and controls per gene as test statistic, and calculated the probability of observing a test statistic by chance from 10,000 permutations. Multiple testing was empirically corrected using ‘minP’ procedure^54,100^. See Supplementary Information for details.

### Data and Code availability

All data presented in this study are accessible under BioProject ID: PRJNA305296. The code is available through R package Rplinkseq and PLINK1.9.

## AUTHOR CONTRIBUTIONS

Conceived and designed the experiments: KLT, EKK, GF, HJN, RT; Analyzed the data: HJN, RT, EKK, JF, CO’D, RS, DH, DPG, KLT; Wrote the paper: HJN, DPG, EKK, KLT; Performed sequence capture: RT, HJN; Performed EMSA: RS; Diagnosed/collected samples: MW, H-JG, SR, CAM, SES, SAR, MAJ, JAK, CR, EG, GLH, DC, EA, SW, PDP, CH, MP, CP; Coordinated/ prepared samples and data generation: JJ, MK, GVG

## ACKNOWLEDGEMENTS

We thank the participating individuals for their support, Eric S. Lander, Steven E. Hyman, Jessica Alföldi and Kaitlin Samocha for valuable inputs, Leslie Gaffney for help with illustrations, Jeremiah M. Scharf for sample contribution and discussions, and Broad Genomics Platform for sample processing, sequencing and genotyping. HJN is supported by AKC Health Foundation, CR by Swedish Research Council (K2013-61P-22168), KLT by Swedish Medical Research Council and European Research Council and EKK by NIH NIMH (1R21MH109938-01). A Broad Institute SPARC grant supported this work.

## References

1. Pauls, D. L. The genetics of obsessive-compulsive disorder: a review. Dialogues Clin. Neurosci. 12, 149–163 (2010).

2. Stewart, S. E. et al. Genome-wide association study of obsessive-compulsive disorder. Mol. Psychiatry 18, 788–798 (2013).

3. Mattheisen, M. et al. Genome-wide association study in obsessive-compulsive disorder: results from the OCGAS. Mol. Psychiatry (2014). doi:10.1038/mp.2014.43.

4. Bienvenu, O. J. et al. Sapap3 and pathological grooming in humans: Results from the OCD collaborative genetics study. Am. J. Med. Genet. B Neuropsychiatr. Genet. 150B, 710–720 (2009).

5. Karagiannidis, I. et al. Replication of association between a SLITRK1 haplotype and Tourette Syndrome in a large sample of families. Mol. Psychiatry 17, 665–668 (2012).

6. Welch, J. M. et al. Cortico-striatal synaptic defects and OCD-like behaviours in Sapap3-mutant mice. Nature 448, 894–900 (2007).

7. Ahmari, S. E. et al. Repeated cortico-striatal stimulation generates persistent OCD-like behavior. Science 340, 1234–1239 (2013).

8. Saxena, S. & Rauch, S. L. Functional neuroimaging and the neuroanatomy of obsessive-compulsive disorder. Psychiatr. Clin. North Am. 23, 563–586 (2000).

9. Karlsson, E. K. & Lindblad-Toh, K. Leader of the pack: gene mapping in dogs and other model organisms. Nat. Rev. Genet. 9, 713–725 (2008).

10. Tang, R. et al. Candidate genes and functional noncoding variants identified in a canine model of obsessive-compulsive disorder. Genome Biol. 15, R25 (2014).

11. Dodman, N. H. et al. A canine chromosome 7 locus confers compulsive disorder susceptibility. Mol. Psychiatry 15, 8–10 (2010).

12. Overall, K. L. Natural animal models of human psychiatric conditions: assessment of mechanism and validity. Prog. Neuropsychopharmacol. Biol. Psychiatry 24, 727–776 (2000).

13. Geschwind, D. H. & Flint, J. Genetics and genomics of psychiatric disease. Science 349, 1489–1494 (2015).

14. Gratten, J., Wray, N. R., Keller, M. C. & Visscher, P. M. Large-scale genomics unveils the genetic architecture of psychiatric disorders. Nat. Neurosci. 17, 782–790 (2014).

15. Ting, J. T. & Feng, G. Neurobiology of obsessive-compulsive disorder: insights into neural circuitry dysfunction through mouse genetics. Curr. Opin. Neurobiol. 21, 842–848 (2011).

16. Sullivan, P. F., Daly, M. J. & O’Donovan, M. Genetic architectures of psychiatric disorders: the emerging picture and its implications. Nat. Rev. Genet. 13, 537–551 (2012).

17. Ioannidis, J. P. A. Commentary: grading the credibility of molecular evidence for complex diseases. Int. J. Epidemiol. 35, 572–8; discussion 593–6 (2006).

18. Farrell, M. S. et al. Evaluating historical candidate genes for schizophrenia. Mol. Psychiatry 20, 555–562 (2015).

19. Sobrin, L. et al. Candidate gene association study for diabetic retinopathy in persons with type 2 diabetes: the Candidate gene Association Resource (CARe). Invest. Ophthalmol. Vis. Sci. 52, 7593–7602 (2011).

20. Fuchsberger, C. et al. The genetic architecture of type 2 diabetes. Nature (2016). doi:10.1038/mp.2014.43

21. D’Gama, A. M. et al. Targeted DNA Sequencing from Autism Spectrum Disorder Brains Implicates Multiple Genetic Mechanisms. Neuron 88, 910–917 (2015).

22. Bertram, L. & Tanzi, R. E. Thirty years of Alzheimer’s disease genetics: the implications of systematic meta-analyses. Nat. Rev. Neurosci. 9, 768–778 (2008).

23. Cohen, J. et al. Low LDL cholesterol in individuals of African descent resulting from frequent nonsense mutations in PCSK9. Nat. Genet. 37, 161–165 (2005).

24. Cohen, J. C., Boerwinkle, E., Mosley, T. H., Jr. & Hobbs, H. H. Sequence variations in PCSK9, low LDL, and protection against coronary heart disease. N. Engl. J. Med. 354, 1264–1272 (2006).

25. Rivas, M. A. et al. Deep resequencing of GWAS loci identifies independent rare variants associated with inflammatory bowel disease. Nat. Genet. 43, 1066–1073 (2011).

26. Jallow, M. et al. Genome-wide and fine-resolution association analysis of malaria in West Africa. Nat. Genet. 41, 657–665 (2009).

27. Gutierrez-Achury, J. et al. Fine mapping in the MHC region accounts for 18% additional genetic risk for celiac disease. Nat. Genet. 47, 577–578 (2015).

28. Roth, E. M., McKenney, J. M., Hanotin, C., Asset, G. & Stein, E. A. Atorvastatin with or without an antibody to PCSK9 in primary hypercholesterolemia. N. Engl. J. Med. 367, 1891–1900 (2012).

29. Stein, E. A. et al. Effect of a monoclonal antibody to PCSK9 on LDL cholesterol. N. Engl. J. Med. 366, 1108–1118 (2012).

30. Warr, A. et al. Exome Sequencing: Current and Future Perspectives. G3 5, 1543–1550 (2015).

31. Gusev, A. et al. Partitioning heritability of regulatory and cell-type-specific variants across 11 common diseases. Am. J. Hum. Genet. 95, 535–552 (2014).

32. Finucane, H. K. et al. Partitioning heritability by functional annotation using genome-wide association summary statistics. Nat. Genet. 47, 1228–1235 (2015).

33. Lek, M. et al. Analysis of protein-coding genetic variation in 60,706 humans. Nature 536, 285–291 (2016).

34. Heiman, M. et al. A translational profiling approach for the molecular characterization of CNS cell types. Cell 135, 738–748 (2008).

35. Collins, M. O. et al. Molecular characterization and comparison of the components and multiprotein complexes in the postsynaptic proteome. J. Neurochem. 97 Suppl 1, 16–23 (2006).

36. Peng, J. et al. Semiquantitative proteomic analysis of rat forebrain postsynaptic density fractions by mass spectrometry. J. Biol. Chem. 279, 21003–21011 (2004).

37. Lein, E. S. et al. Genome-wide atlas of gene expression in the adult mouse brain. Nature 445, 168–176 (2007).

38. Basu, S. N., Kollu, R. & Banerjee-Basu, S. AutDB: a gene reference resource for autism research. Nucleic Acids Res. 37, D832–6 (2009).

39. Bassett, A. S., Marshall, C. R., Lionel, A. C., Chow, E. W. & Scherer, S. W. Copy number variations and risk for schizophrenia in 22q11.2 deletion syndrome. Hum. Mol. Genet. 17, 4045–4053 (2008).

40. McCarthy, S. E. et al. Microduplications of 16p11.2 are associated with schizophrenia. Nat. Genet. 41, 1223–1227 (2009).

41. Kumar, R. A. et al. Recurrent 16p11.2 microdeletions in autism. Hum. Mol. Genet. 17, 628–638 (2008).

42. Stefansson, H. et al. Large recurrent microdeletions associated with schizophrenia. Nature 455, 232–236 (2008).

43. van der Zwaag, B. et al. A co-segregating microduplication of chromosome 15q11.2 pinpoints two risk genes for autism spectrum disorder. Am. J. Med. Genet. B Neuropsychiatr. Genet. 153B, 960–966 (2010).

44. Tabares-Seisdedos, R. & Rubenstein, J. L. Chromosome 8p as a potential hub for developmental neuropsychiatric disorders: implications for schizophrenia, autism and cancer. Mol. Psychiatry 14, 563–589 (2009).

45. Shugart, Y. Y. et al. Genomewide linkage scan for obsessive-compulsive disorder: evidence for susceptibility loci on chromosomes 3q, 7p, 1q, 15q, and 6q. Mol. Psychiatry 11, 763–770 (2006).

46. Lindblad-Toh, K. et al. A high-resolution map of human evolutionary constraint using 29 mammals. Nature 478, 476–482 (2011).

47. Wei, Z., Wang, W., Hu, P., Lyon, G. J. & Hakonarson, H. SNVer: a statistical tool for variant calling in analysis of pooled or individual next-generation sequencing data. Nucleic Acids Res. 39, e132 (2011).

48. Thurman, R. E. et al. The accessible chromatin landscape of the human genome. Nature 489, 75–82 (2012).

49. Pruitt, K. D. et al.RefSeq: an update on mammalian reference sequences. Nucleic Acids Res. 42, D756–63 (2014).

50. Wang, K., Li, M. & Hakonarson, H. ANNOVAR: functional annotation of genetic variants from high-throughput sequencing data. Nucleic Acids Res. 38, e164 (2010).

51. Dunham, I. et al. An integrated encyclopedia of DNA elements in the human genome. Nature 489, 57–74 (2012).

52. Davydov, E. V. et al. Identifying a high fraction of the human genome to be under selective constraint using GERP++. PLoS Comput. Biol. 6, e1001025 (2010).

53. The 1000 Genomes Project Consortium. A global reference for human genetic variation. Nature 526, 68–74 (2015).

54. Purcell, S. M. et al. A polygenic burden of rare disruptive mutations in schizophrenia. Nature 506, 185–190 (2014).

55. Kiezun, A. et al. Exome sequencing and the genetic basis of complex traits. Nat. Genet. 44, 623–630 (2012).

56. NCBI Resource Coordinators. Database resources of the National Center for Biotechnology Information. Nucleic Acids Res. 44, D7–D19 (2016).

57. Porcelli, S. et al. Pharmacogenetics of antidepressant response. J. Psychiatry Neurosci. 36, 87–113 (2011).

58. Chen, Y.-K., Y.-K., C. & Y.-P., H. Cortactin-Binding Protein 2 Modulates the Mobility of Cortactin and Regulates Dendritic Spine Formation and Maintenance. Journal of Neuroscience 32, 1043–1055 (2012).

59. Castermans, D. et al. Identification and characterization of the TRIP8 and REEP3 genes on chromosome 10q21.3 as novel candidate genes for autism. Eur. J. Hum. Genet. 15, 422–431 (2007).

60. Roadmap Epigenomics Consortium et al. Integrative analysis of 111 reference human epigenomes. Nature 518, 317–330 (2015).

61. Pauls, D. L., Abramovitch, A., Rauch, S. L. & Geller, D. A. Obsessive-compulsive disorder: an integrative genetic and neurobiological perspective. Nat. Rev. Neurosci. 15, 410–424 (2014).

62. Kheradpour, P. et al. Systematic dissection of regulatory motifs in 2000 predicted human enhancers using a massively parallel reporter assay. Genome Res. 23, 800–811(2013).

63. Becamel, C. et al. The serotonin 5-HT2A and 5-HT2C receptors interact with specific sets of PDZ proteins. J. Biol. Chem. 279, 20257–20266 (2004).

64. Schneider, J. W. Caveats for using statistical significance tests in research assessments. J. Informetr. 7, 50–62 (2013/1).

65. Ebert, D. H. & Greenberg, M. E. Activity-dependent neuronal signalling and autism spectrum disorder. Nature 493, 327–337 (2013).

66. Barak, B. & Feng, G. Neurobiology of social behavior abnormalities in autism and Williams syndrome. Nat. Neurosci. 19, 647–655 (2016).

67. Park, J.-H. et al. Distribution of allele frequencies and effect sizes and their interrelationships for common genetic susceptibility variants. Proc. Natl. Acad. Sci. U. S. A. 108, 18026–18031 (2011).

68. Huang, J. et al. Improved imputation of low-frequency and rare variants using the UK10K haplotype reference panel. Nat. Commun. 6, 8111 (2015).

69. The UK10K Consortium. The UK10K project identifies rare variants in health and disease. Nature 526, 82–90 (2015).

70. Wu, M. C. et al. Rare-variant association testing for sequencing data with the sequence kernel association test. Am. J. Hum. Genet. 89, 82–93 (2011).

71. Adzhubei, I. A. et al. A method and server for predicting damaging missense mutations. Nat. Methods 7, 248–249 (2010).

72. Zuk, O. et al. Searching for missing heritability: designing rare variant association studies. Proc. Natl. Acad. Sci. U. S. A. 111,E455–64 (2014).

73. de Wit, J. et al. LRRTM2 interacts with Neurexin1 and regulates excitatory synapse formation. Neuron 64, 799–806 (2009).

74. Surmeier, D. J., Ding, J., Day, M., Wang, Z. & Shen, W. D1 and D2 dopamine-receptor modulation of striatal glutamatergic signaling in striatal medium spiny neurons. Trends Neurosci. 30, 228–235 (2007).

75. Sudhof, T. C. Neuroligins and neurexins link synaptic function to cognitive disease. Nature 455, 903–911 (2008).

76. Jenkins, A. K. et al. Neurexin 1 (NRXN1) splice isoform expression during human neocortical development and aging. Mol. Psychiatry 21, 701–706 (2016).

77. Rujescu, D. et al. Disruption of the neurexin 1 gene is associated with schizophrenia. Hum. Mol. Genet. 18, 988–996 (2009).

78. Blackstone, C., O’Kane, C. J. & Reid, E. Hereditary spastic paraplegias: membrane traffic and the motor pathway. Nat. Rev. Neurosci. 12, 31–42 (2011).

79. Kala, K. et al. Gata2 is a tissue-specific post-mitotic selector gene for midbrain GABAergic neurons. Development 136, 253–262 (2009).

80. Ling, B. E.Obsessive compulsive disorder research. (Nova Science Publishers, 2005).

81. Chen, Y. K. & Hsueh, Y. P. Cortactin-binding protein 2 modulates the mobility of cortactin and regulates dendritic spine formation and maintenance. J. Neurosci. 32, 1043–1055 (2012).

82. El Sayegh, T. Y. et al. Cortactin associates with N-cadherin adhesions and mediates intercellular adhesion strengthening in fibroblasts. J. Cell Sci. 117, 5117–5131 (2004).

83. Taylor, S. Molecular genetics of obsessive-compulsive disorder: a comprehensive meta-analysis of genetic association studies. Mol. Psychiatry 18, 799–805 (2013).

84. Dodman, N. H. et al. Genomic Risk for Severe Canine Compulsive Disorder, a Dog Model of Human OCD. Int. J. Appl. Res. Vet. Med. 14, (2016).

85. Edlund, C. K., Conti, D. V. & Van Den Berg, D. J. rAggr. (2016). Available at: http://raggr.usc.edu. (Accessed: 8th August 2016)

86. Shmelkov, S. V. et al. Slitrk5 deficiency impairs corticostriatal circuitry and leads to obsessive-compulsive-like behaviors in mice. Nat. Med. 16, 598–602, 1p following 602 (2010).

87. Graf, E. R., Zhang, X., Jin, S. X., Linhoff, M. W. & Craig, A. M. Neurexins induce differentiation of GABA and glutamate postsynaptic specializations via neuroligins. Cell 119, 1013–1026 (2004).

88. Doly, S. & Marullo, S. Gatekeepers Controlling GPCR Export and Function. Trends Pharmacol. Sci. 36, 636–644 (2015).

89. Marin, O. & Rubenstein, J. L. A long, remarkable journey: tangential migration in the telencephalon. Nat. Rev. Neurosci. 2, 780–790 (2001).

90. McMahon, F. J. et al. Variation in the gene encoding the serotonin 2A receptor is associated with outcome of antidepressant treatment. Am. J. Hum. Genet. 78, 804–814 (2006).

91. Noordam, R. et al. Identifying genetic loci associated with antidepressant drug response with drug–gene interaction models in a population-based study. J. Psychiatr. Res. 62, 31–37 (2015/3).

92. Schizophrenia Working Group of the Psychiatric Genomics Consortium. Biological insights from 108 schizophrenia-associated genetic loci. Nature 511, 421–427 (2014).

93. Schwarz, D. S. & Blower, M. D. The endoplasmic reticulum: structure, function and response to cellular signaling. Cell. Mol. Life Sci. 73, 79–94 (2016).

94. The UniProt Consortium. UniProtKB-P28223 (5HT2A_HUMAN). (2016). Available at: http://www.uniprot.org/uniprot/P28223. (Accessed: 8th August 2016.

95. Ashley, E. A. Towards precision medicine. Nat. Rev. Genet. 17, 507–522 (2016).

96. Popejoy, A. B. & Fullerton, S. M. Genomics is failing on diversity. Nature 538, 161–164 (2016).

97. Bustamante, C. D., Burchard, E. G. & De la Vega, F. M. Genomics for the world. Nature 475, 163–165 (2011).

98. American Psychiatric Association. Diagnostic and statistical manual of mental disorders : DSM-IV-TR. (American Psychiatric Association, 2000).

99. Chang, C. C. et al. Second-generation PLINK: rising to the challenge of larger and richer datasets. Gigascience 4, 7 (2015).

100. Kiezun, A. et al. Exome sequencing and the genetic basis of complex traits. Nat. Genet. 44, 623–630 (2012).

101. Lambe, E. K., Fillman, S. G., Webster, M. J. & Shannon Weickert, C. Serotonin receptor expression in human prefrontal cortex: balancing excitation and inhibition across postnatal development. PLoS One 6, e22799 (2011).

102. Chen, Y. K., Chen, C. Y. & Hu, H. T. CTTNBP2, but not CTTNBP2NL, regulates dendritic spinogenesis and synaptic distribution of the striatin–PP2A complex. Mol. Biol. Cell (2012).

103. Niknafs, N. et al. MuPIT interactive: webserver for mapping variant positions to annotated, interactive 3D structures. Hum. Genet. 132, 1235–1243 (2013).

104. Chen, F., Venugopal, V., Murray, B. & Rudenko, G. The structure of neurexin 1α reveals features promoting a role as synaptic organizer. Structure 19, 779–789 (2011).

